# Reforestation under climate change - a cross-regional deadwood retention experiment to develop multifunctional forests on disturbed Norway spruce areas

**DOI:** 10.64898/2026.01.12.698939

**Authors:** Birgitta Putzenlechner, Martin Zwanzig, Claus Bässler, Markus Bernhardt-Römermann, Janina Ebert, Simon Grieger, Philipp Koal, Henrik Oechler, Dorothea Peter, Ingolf Profft, Sascha Spalek, Florian Steinebrunner, Alexander Tischer, Simon Thorn, Alexandra Wehnert, Xinying Zhou, Franka Huth

## Abstract

The impact of extreme weather events such as droughts, heatwaves, storms and the associated intensification of bark beetle outbreaks in Norway spruce forests in recent years, has led to a large-scale forest loss in Central Europe. There are various options for post-disturbance management, from complete clearing to no intervention, but their pros and cons – in the context of multifunctional forest management – are not resolved. The “ResEt-Fi” project (“Pioneering reforestation: Regional management for establishing multifunctional forests on post-disturbed Norway spruce areas”) consortium implemented a cross-regional field trial to test how specific practices of deadwood retention affect the ecological conditions and subsequent reforestation dynamics of natural regeneration and planted trees. It aims at generating climate-stable mixed forests within an ecological valuable transition, balancing ecosystem functioning and socio-economic benefits. As an outcome for stakeholders and forest practitioners, guidelines and showcase scenarios for facilitating the selection of management procedures will be created. In this article, we provide insight into (i) the applied post-disturbance deadwood management practices and the rationale for their selection, (ii) the experimental setup of our cross-regional field trial, (iii) the initial post-disturbance status and perspective with respect to regional site conditions and deadwood treatment. While the setup and site conditions will primarily serve as a reference for subsequent studies in our joint project, this is the first systematic, cross-regional study on the effects of the management options standing deadwood, high stumps and clearing on microclimate, biodiversity, and tree regeneration in the critical early stages of forest regeneration after disturbance.

## 1. Introduction

In recent years, forest ecosystems across Central Europe have been increasingly affected by climate-induced disturbances (Forzieri et al., 2021; Knutzen et al., 2025; Patacca et al., 2023). The combination of prolonged droughts, heat extremes, and subsequent bark beetle outbreaks have triggered widespread forest disturbances and dieback, highlighting the vulnerability of current forest landscapes to compound stressors (Hartmann et al., 2025; Senf, Buras, Zang, Rammig, & Seidl, 2020; Senf & Seidl, 2021). Although tree species, previously considered well-adapted to the Central European climate, like European beech (*Fagus sylvatica* L.), are affected as well (Leuschner, 2020; West, Morley P. J., Sump, & Donoghue, 2022), particularly severe and large-scale impacts have been observed for Norway spruce (*Picea abies* (L.) H. Karst.) (Hlásny et al., 2021; Obladen et al., 2021). Until today, Norway spruce as even-aged and mono-layered stands with low proportions of mixed tree species dominate large areas. This situation is a result of historical disturbances, reparation cuts, and silvicultural practices of thinning from below, driven by the motivation to maximize timber production of one tree species per forest area (Nyland, 2016; Spiecker et al., 2004). Although the vulnerability of this species to wind breaks and more frequent droughts under a warming climate has been emphasized in numerous studies (Anders et al., 2025; Buras & Menzel, 2019; Kunert et al., 2022; Lévesque et al., 2013), the intensity of disturbance in recent years was unprecedented (Hartmann et al., 2025). The high temporal dynamics of disturbances have forced forest management to act swiftly, often in an evidence-limited space where ecological and socio-economic outcomes remain uncertain.

Based on nation-wide remote sensing-based assessment, a forest canopy cover loss of more than 500,000 ha has been reported across Germany (Thonfeld et al., 2022), mainly due to bark beetle outbreaks. Between 2018 and early 2025, approximately 308 million solid cubic meters of calamity wood, resulting from storm damage, drought, or pest infestation, were recorded in Germany (Bundesministerium für Landwirtschaft, Ernährung und Heimat (BMLEH), 2025), which includes more than 282 million m³ of coniferous and 26 million m³ of deciduous wood. The volume harvested represents over 20% of the national Norway spruce stock reported in the 2012 Federal Forest Inventory, indicating a significant loss. Major damage has occurred particularly in forests of low mountain regions. Reforestation needs exceed 500,000 hectares nationwide (Bundesministerium für Landwirtschaft, Ernährung und Heimat (BMLEH), 2025) and the estimated economic loss to the forestry sector from 2018 to 2020 surpasses €12.7 billion (Möhring et al., 2021). In Thuringia, a German federal state with a regionally important timber industry, as of March 2023, approximately 86,000 hectares were affected. This means that nearly 20% of the state’s forest cover was lost within six years; in March 2025 the amount of loss in the forest area has increased to approximately 133,000 hectares (ThüringenForst AöR, 2023; ThüringenForst AöR, 2025). In contrast to past storm-related, rather temporary forest disturbances, the current situation represents a new societal challenge, as severe consequences are expected for the diverse ecosystem services and functions provided by forests, with implications for human well-being and the structure of cultural landscapes (Brockerhoff et al., 2017; Morris et al., 2018). Based on the study by Fleischer et al. (2017) in the Norway spruce forests of the Tatra Mountains (Slovakia), it can be assumed that cultural (e.g., number of visitors, attractiveness) and provisioning (e.g., gross and net primary production) services decline within the first ten years after the disturbance, while regulating services (e.g., respiration, soil moisture) stabilize more quickly.

Common management responses to bark beetle-related disturbances frequently involve sanitary and salvage logging, both including the complete removal of dead and dying trees (Leverkus, Lindenmayer, Thorn, & Gustafsson, 2018). While phytosanitary goals are the motivation for sanitary logging, such as limiting the further spread of insect infestations, economic incentives to recover the residual timber value associated with salvage logging are also a major driving force. The commercial worth of deadwood, particularly standing trunks, encourages salvage logging with extensive clearing of disturbed areas, often resulting in the near-total removal of remaining forest structural elements (Keenan & Kimmins, 1993). In practice, both approaches often occur simultaneously, as the rapid spread of bark beetle infestations necessitate prompt and extensive interventions; therefore, we refer to these combined practices as salvage logging in the following. In many cases, clear-cutting is carried out to facilitate site preparation and subsequent replanting, sometimes preceded by mulching or other forms of biomass reduction. The large-scale clearings of calamities occurring over extensive areas, have the potential to fundamentally alter forest structure and ecosystem functions (Chianucci et al., 2024). Specifically, such interventions may lead to ecological consequences like the loss of microclimatic buffering, declines in habitat heterogeneity and biodiversity, soil degradation, or impaired ecosystem resilience over the long term (Leverkus et al., 2021b; Thorn et al., 2018). However, to meet future demands for multifunctional forest use under changing climatic conditions, the development of stable, resilient, and vital forest ecosystems on those disturbed areas is urgently needed.

The close relationship between forest structure and microclimatic conditions is well known (Atkins, Shiklomanov, Mathes, Bond-Lamberty, & Gough, 2023; Frenne et al., 2021). Large-scale canopy loss following disturbance leads to strong microclimatic amplification, characterized by increased daytime temperatures and intensified nocturnal cooling. This disrupts or even eliminates the buffered forest interior climate that typically mitigates temperature and moisture extremes, with potentially negative consequences for natural regeneration and reforestation success, particularly under extensive disturbance (Hannerz & Hånell, 1997). At the same time, the widespread removal of deadwood further amplifies these effects by eliminating essential structural and resource components that support roughly one-third of all forest-dwelling species (Siitonen, 2001). The combined loss of microclimatic buffering and habitat structures not only reduces biodiversity but also impairs key processes that facilitate the establishment of forest regeneration. Importantly, remaining deadwood structures, such as standing snags and downed logs, can mitigate these stresses by providing ground-level shade, reducing wind exposure, and limiting soil evaporation, thereby alleviating heat and water stress for establishing seedlings and saplings (Steinebrunner, Tischer, Medicus, Huth, & Bernhardt-Römermann, 2025).

From an ecological perspective, the retention of deadwood contributes significantly to structural heterogeneity and nutrient cycling, thereby expanding habitat availability and supporting diverse food webs (Müller et al., 2023; Thorn et al., 2018). As deadwood decomposes into coarse and fine woody debris, it plays a key role in the formation of organic soil layers, enhancing water retention capacity and nutrient availability, both critical for seedling survival and early growth (Laiho & Prescott, 2004). Importantly, deadwood is a keystone habitat for saproxylic organisms, including insects (e.g. beetles), fungi, mosses, and lichens, many of which depend on specific stages of wood decay. These species contribute to decomposition processes, nutrient mobilization, and soil formation, and several also play roles in natural pest regulation (Seibold et al., 2016). Thus, a high diversity of these organisms may enhance functional stability during the regeneration phase, as suggested by the insurance hypothesis of Yachi and Loreau (1999), through supporting processes such as litter decomposition and mycorrhizal networks.

Overall, as salvage logging in response to forest disturbances, such as the wide-spread bark beetle infestations in Norway spruce, will become an increasingly prevalent second disturbance regime in Central Europe, its ecological implications warrant critical evaluation and assessment of alternative management strategies while addressing economic considerations (Knoke et al., 2021). Post-disturbance management strategies commonly rely on either salvage logging or full preservation of unlogged forest stands, while structured intermediate approaches with temporal and spatial coordination remain largely absent in forestry practice. Following the concept of retention forestry, to retain structural and compositional heterogeneity of both remaining and dead trees (Gustafsson et al., 2012), post-disturbance management strategies that retain structural elements and promote microclimatic buffering could play a crucial role in supporting the successful establishment and survival of forest regeneration (Leverkus et al., 2021b; Storch et al., 2020; Zhang, Sjögren, & Jönsson, 2024). This deadwood-induced amelioration could facilitate survival and growth rates of both planted seedlings and natural regeneration, ultimately increasing the speed of reforestation of disturbed sites and their transition into economically productive forest sites (Castro et al., 2011; Marangon, Marchi, & Lingua, 2022; Swanson, Magee, Nelson, Engstrom, & Adams, 2023). Thus, either removing none or only a limited portion of dead trunks could therefore be advantageous from both the ecological and economic perspective.

Research on retention forestry and the effects of structural diversity has largely concentrated on the management of vital stands (Huang et al., 2023; Menge, Magdon, Wöllauer, & Ehbrecht, 2023; Nirhamo, Hämäläinen, Junninen, & Kouki, 2023; Storch et al., 2020; Zhang et al., 2024) or in the context of wildfires (Castro et al., 2011; Lingua et al., 2023; Marshall et al., 2023; Marzano, Garbarino, Marcolin, Pividori, & Lingua, 2013), while post-disturbance management strategies in bark-beetle-induced die-back of Norway spruce —beyond full retention of deadwood trees (Schroeder, 2007)—remain understudied in Central Europe. Although some recent studies have investigated retaining management strategies, examining how different types and amounts of biological legacies (e.g., downed woody debris to unlogged deadwood patches) affect biodiversity and forest regeneration (Marzano et al., 2013; Swanson et al., 2023), there is a lack of interdisciplinary research and practical benchmarks concerning both the composition and volume of retained structural legacies, particularly regarding the effects of partial deadwood retention following large-scale dieback events (Steinebrunner et al., 2025). Moreover, recent findings from small-scale plot studies (Putzenlechner, Grieger, Czech, & Koal, 2025) need to be scaled up to the landscape level using remote sensing and spatial modeling, in order to capture spatial heterogeneity and to inform regional management planning. While recent studies have highlighted the strong relationship between land use, land cover, and land surface temperatures (Gohr, Blumröder, Sheil, & Ibisch, 2021; Zimmermann, Kruber, Nendel, Munack, & Hildmann, 2024), the implications of these patterns for forest regeneration at landscape-level remain insufficiently explored. In this regard, the consistently elevated summer surface temperatures observed in disturbed areas (Grieger et al., 2025; Hesslerová, Huryna, Pokorný, & Procházka, 2018) underscore the need to further investigate the long-term climatic and ecological feedbacks of different post-disturbance management regimes. The current knowledge gaps limit our understanding of the microclimatic and ecological consequences of nuanced silvicultural interventions on disturbed sites.

At the same time, an effective knowledge transfer strategy is needed to inform practitioners, who urgently require and request spatially explicit guidance under changing climate conditions. Despite increasing recognition of climate-induced forest disturbances among forest practitioners, substantial gaps remain in translating ecological insights into guidelines and best-practice protocols (Nikinmaa et al., 2024). Forestry operations and decisions are constrained by staff shortages, particularly when multiple large-scale disturbance events require rapid intervention, while the market faces challenges in processing and storing large volumes of salvaged timber, due to limited infrastructure, oversupply, and volatile prices (Puettmann et al., 2015). Moreover, forest management operates within a field of tension between ecologically sound measures and their public acceptance, underscoring the necessity of clear, evidence-based guidelines to ensure informed and legitimate decision-making. As such, silvicultural strategies for reforesting affected areas under changing climatic conditions remain under debate, especially given uncertainties in species selection for forest conversion (Albert, Nagel, Nuske, Sutmöller, & Spellmann, 2017). These are further compounded by limited seed availability, scarce operational experience, and difficulties in managing extensive deadwood legacies. Together, these factors highlight the urgent need for integrating ecological research with practical forestry to develop both adaptive and scalable management approaches. Targeted forestry concepts must be developed, particularly regarding ensuring diverse or specific ecosystem services in the future.

Against this background, it becomes clear that an integrated and interdisciplinary assessment is required to jointly evaluate the effects of management measures on abiotic, soil– and biodiversity-related factors in parallel and to identify alternative pathways to complete clearings, focusing on the reforestation chances to consider also the economic point of view. Therefore, local factors such as soil properties, seed tree availability, accessibility and objectives of local forest management must be studied to identify optimal customizations of management strategies.

### 1.1 Research agenda for an integrated assessment of post-disturbance management options

We hereby present an innovative, interdisciplinary research approach for developing future forest management strategies following large-scale disturbances in former Norway spruce-dominated low mountain ranges of Central Europe. This integrated approach includes research, analysis, knowledge transfer and implementation. The interrelationships are researched and analyzed from natural science, forestry science, and socio-economic perspectives. We aim at developing management strategies to reach multifunctional forests under changing environmental conditions and to examine and assess their impact on various ecosystem services and forest functions. The findings are also prepared for practical application to involve multipliers and various stakeholders in the transfer process (e.g., forestry training, forestry and non-forestry stakeholders). Knowledge transfer at the study areas as local learning locations and “real-world forest laboratories” are of essential importance. The creation of on-site research and demonstration platforms allows for the involvement of local stakeholders as multipliers (e.g., interest groups, see above), also ensuring the integration and implementation of empirically gained knowledge to the regional or cross-regional level. Our overall agenda focuses on the following main objectives:

(I) Interdisciplinary assessment of treatment variants with different deadwood management and comparison to still intact Norway spruce stands along representative regions.
(II) Development of recommendations for action and evaluation schemes for the intensity of salvage logging and the retention of different deadwood structures.
(III) Upscaling of observed local effects and dynamics for recommendations for action ensuring the fulfillment of ecosystem-specific services at larger temporal and spatial scales such as regions or landscapes.

Current forest management in Central Europe’s Norway spruce-dominated low mountain ranges is thus enriched by a holistic approach that could prove effective for the development and establishment of multifunctional and climate-adapted forests.

### 1.2 Research questions and approach

Given the current widespread salvage logging as a response to Norway spruce calamity, and the overall goal to establish multifunctional forests, we focus on the following research questions:

i) Which effects do different post-disturbance management strategies have regarding forest functions and ecosystem services?
ii) Which site-specific management approach at the stand level provides the most versatile development option for climate-resilient and multifunctional future forests?
iii) How can these site-specific insights be translated into regional management and adaptation strategies to address climate change impacts more broadly?

To answer these questions, interdisciplinary efforts as depicted in **Figure 1** are necessary, including aspects of (micro-)climatic effects, soil ecology, vegetation dynamics of the herbaceous layer, shrubs and tree species, effects on microbial communities, birds and wood-dwelling beetles. Moreover, to upscale findings from plot to regional scale, a multisensory remote sensing approach is implemented (spatial upscaling) so that ultimately, predictions of the dynamics of various future scenarios are derived from process-based simulations using the individual-based forest landscape and disturbance model iLand (temporal upscaling).

**Figure 1:**
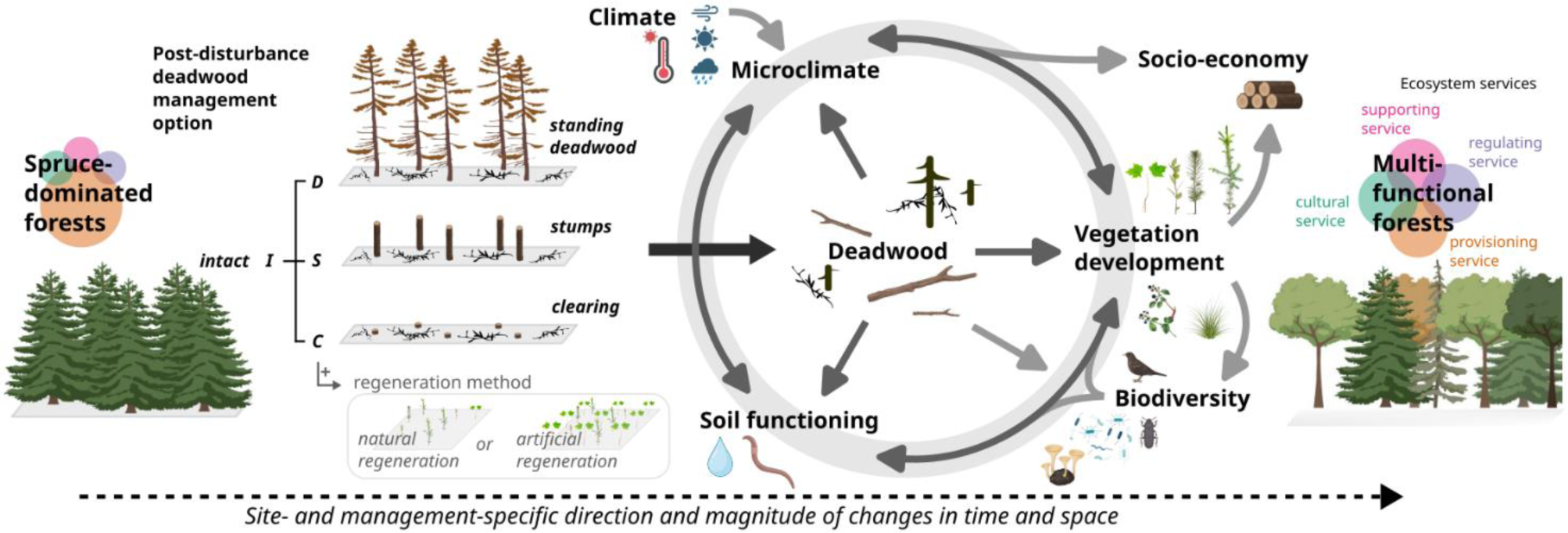
Schematic overview of the effects of post-disturbance deadwood management options on interactions within the disturbed area ecosystem and the paths to developing multifunctional forests in the face of climate change.

It becomes clear that the research activities need to target multi-year monitoring of forest development (>3-5 years). In this way, implemented real-world forest laboratories can provide an opportunity to assess the resilience of ecosystems, also capturing slow-moving changes. With research activities delivering a broad database, established plots can serve as continuous, long-term learning objects for silvicultural and forest ecology training courses, workshops, and excursions for a wide range of stakeholders and partners.

## 2. Experimental design

Research is centered in Thuringia, a federal state in Central Germany, including both disturbed and undisturbed forest stands as current hotspots of bark beetle-related forest disturbance in Germany. Within three model regions, a nested experimental design is applied that allows for assessing the effects of different post-disturbance management from local to regional scale (**Fig. 2**). The local scale comprises plots and sub-plots, enabling detailed characterization of the site-specific conditions of selected disturbance areas and post-disturbance management treatments, supported by intensive empirical and interdisciplinary investigations. The field trial comprises three post-disturbance treatment variants: standing deadwood patches (“D”), high stumps (dead trees cut at 2–3 m height, “S”) and cleared areas (salvage logging, “C”) as well as monitoring of intact Norway spruce stands (“I”).

**Figure 2:**
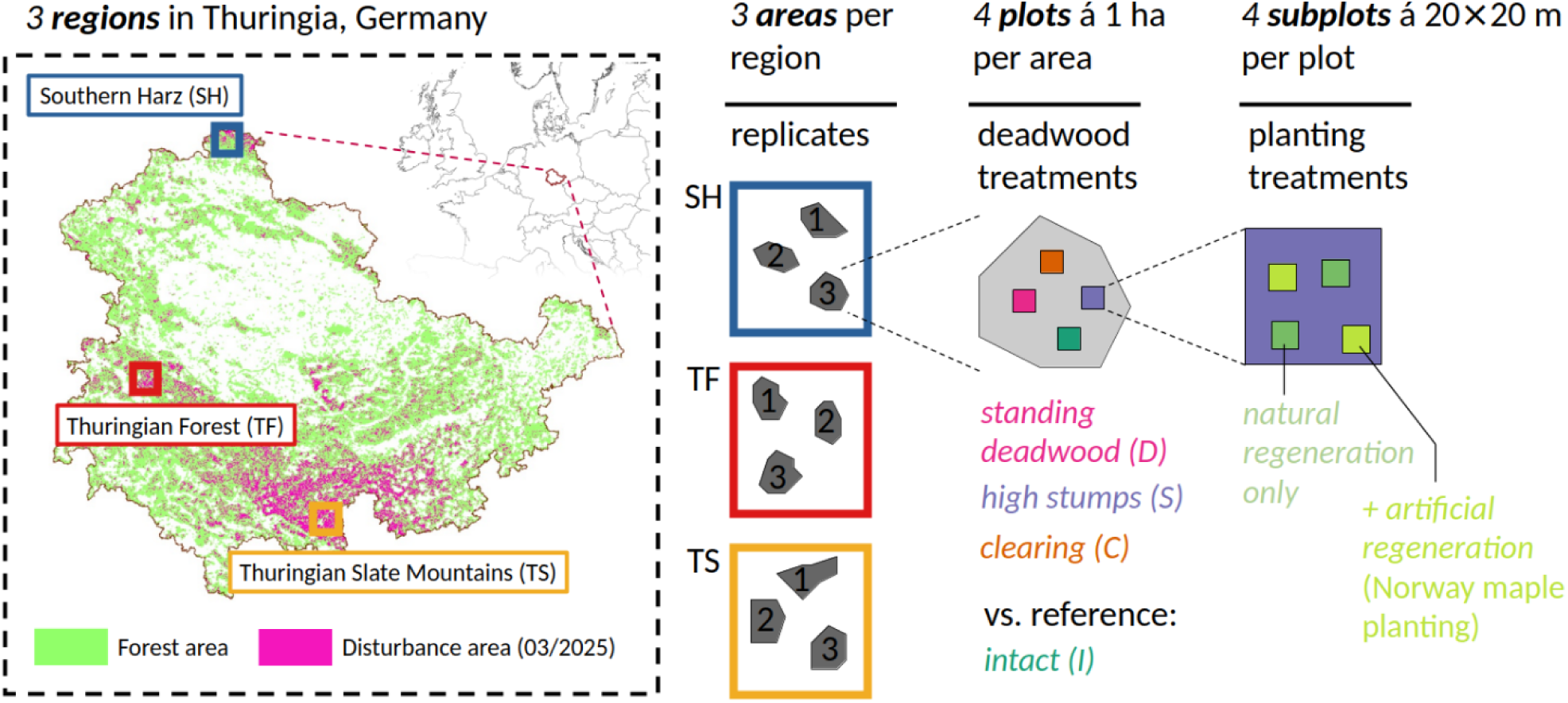
Nested experimental approach with model regions within Thuringia, Germany, areas as replicates within each region (see Table 1 for details), deadwood retention treatment plots within each area and planting treatment subplots (2 of each category, except for intact stands) within each plot.

**Table 1:**
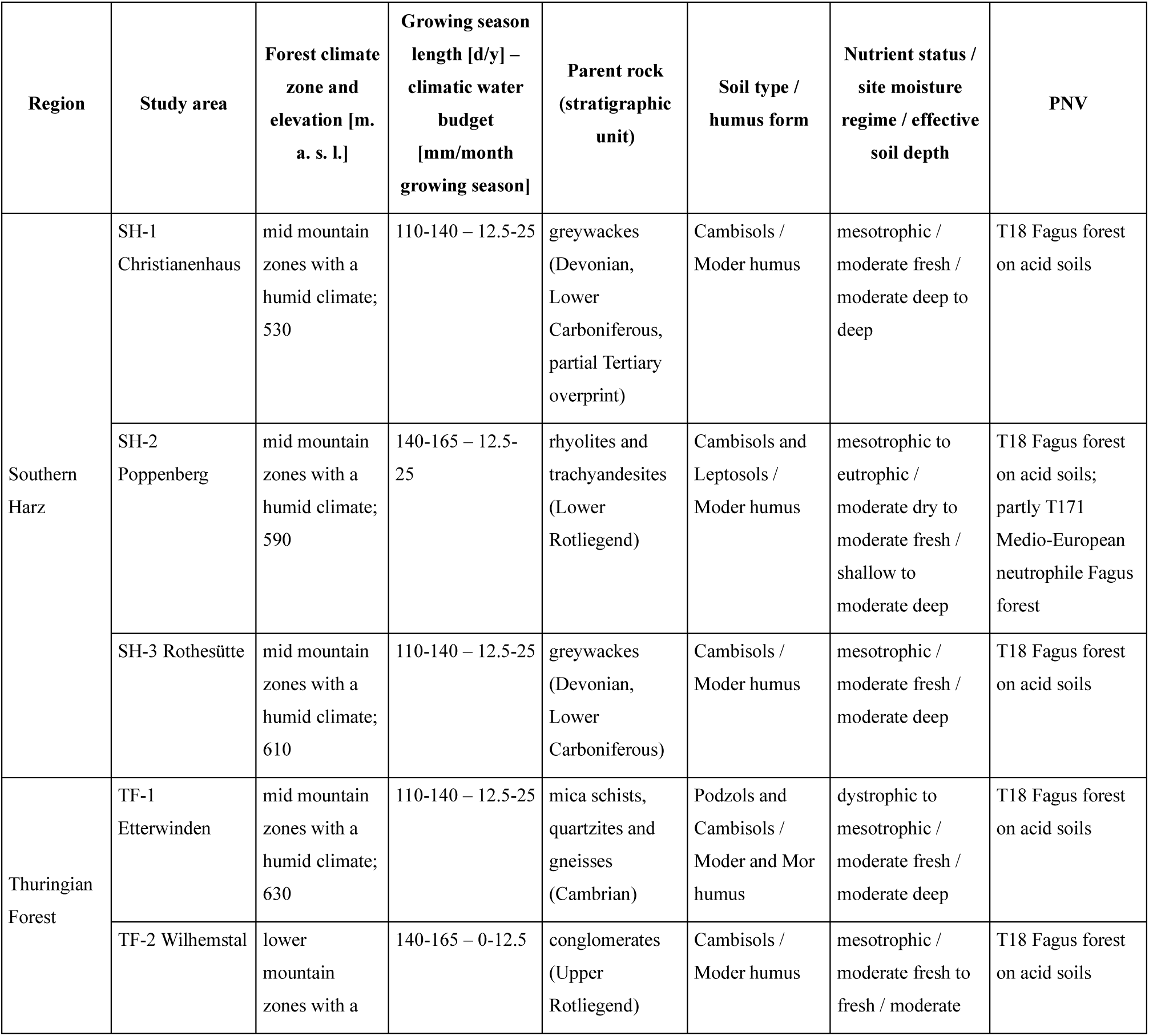
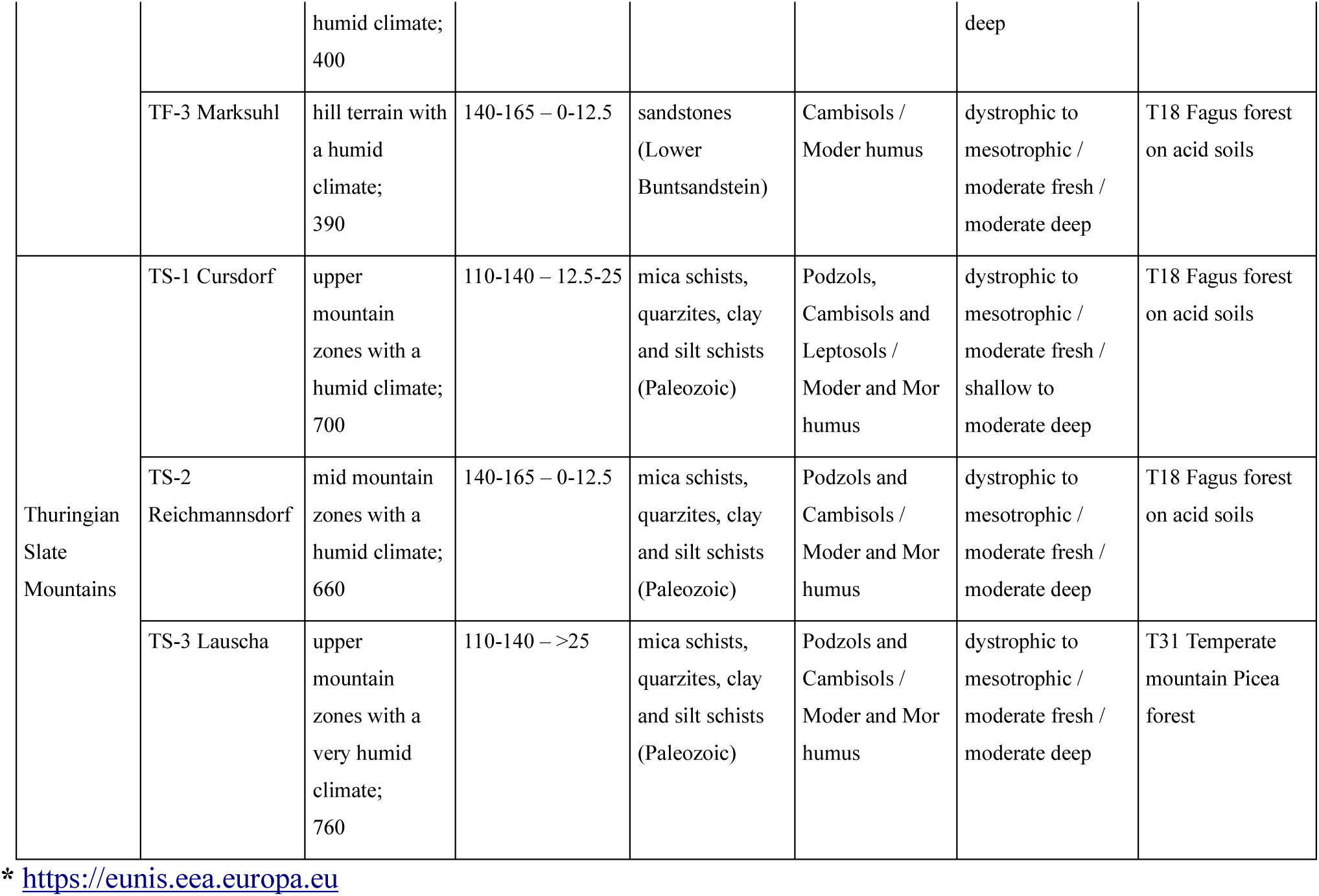
Overview of topographic, climatic and geopedological characteristics of the study areas. Classification of the tree layer potential natural vegetation (PNV) is following the EUNIS classification system of the European Environment Agency*. Across all study areas, existing pre-disturbance stand composition is classified as ‘T3N Coniferous plantation of site-native trees’.

| Region | Study area | Forest climate zone and elevation [m. a. s. l.] | Growing season length [d/y] – climatic water budget [mm/month growing season] | Parent rock (stratigraphic unit) | Soil type / humus form | Nutrient status / site moisture regime / effective soil depth | PNV |
| --- | --- | --- | --- | --- | --- | --- | --- |
| Southern Harz | SH-1 Christianenhaus | mid mountain zones with a humid climate; 530 | 110-140 – 12.5-25 | greywackes (Devonian, Lower Carboniferous, partial Tertiary overprint) | Cambisols / Moder humus | mesotrophic / moderate fresh / moderate deep to deep | T18 Fagus forest on acid soils |
|  | SH-2 Poppenberg | mid mountain zones with a humid climate; 590 | 140-165 – 12.5-25 | rhyolites and trachyandesites (Lower Rotliegend) | Cambisols and Leptosols / Moder humus | mesotrophic to eutrophic / moderate dry to moderate fresh / shallow to moderate deep | T18 Fagus forest on acid soils; partly T171 Medio-European neutrophile Fagus forest |
|  | SH-3 Rothesütte | mid mountain zones with a humid climate; 610 | 110-140 – 12.5-25 | greywackes (Devonian, Lower Carboniferous) | Cambisols / Moder humus | mesotrophic / moderate fresh / moderate deep | T18 Fagus forest on acid soils |
| Thuringian Forest | TF-1 Etterwinden | mid mountain zones with a humid climate; 630 | 110-140 – 12.5-25 | mica schists, quartzites and gneisses (Cambrian) | Podzols and Cambisols / Moder and Mor humus | dystrophic to mesotrophic / moderate fresh / moderate deep | T18 Fagus forest on acid soils |
|  | TF-2 Wilhemstal | lower mountain zones with a humid climate;<br>400 | 140-165 – 0-12.5 | conglomerates (Upper Rotliegend) | Cambisols / Moder humus | mesotrophic / moderate fresh to fresh / moderate deep | T18 Fagus forest on acid soils |
|  | TF-3 Marksuhl | hill terrain with<br>a humid<br>climate;<br>390 | 140-165 – 0-12.5 | sandstones<br>(Lower<br>Buntsandstein) | Cambisols /<br>Moder humus | dystrophic to<br>mesotrophic /<br>moderate fresh /<br>moderate deep | T18 Fagus forest<br>on acid soils |
| Thuringian<br>Slate<br>Mountains | TS-1 Cursdorf | upper<br>mountain<br>zones with a<br>humid climate;<br>700 | 110-140 – 12.5-25 | mica schists,<br>quarzites, clay<br>and silt schists<br>(Paleozoic) | Podzols,<br>Cambisols and<br>Leptosols /<br>Moder and Mor<br>humus | dystrophic to<br>mesotrophic /<br>moderate fresh /<br>shallow to<br>moderate deep | T18 Fagus forest<br>on acid soils |
|  | TS-2<br>Reichmannsdorf | mid mountain<br>zones with a<br>humid climate;<br>660 | 140-165 – 0-12.5 | mica schists,<br>quarzites, clay<br>and silt schists<br>(Paleozoic) | Podzols and<br>Cambisols /<br>Moder and Mor<br>humus | dystrophic to<br>mesotrophic /<br>moderate fresh /<br>moderate deep | T18 Fagus forest<br>on acid soils |
|  | TS-3 Lauscha | upper<br>mountain<br>zones with a<br>very humid<br>climate;<br>760 | 110-140 – >25 | mica schists,<br>quarzites, clay<br>and silt schists<br>(Paleozoic) | Podzols and<br>Cambisols /<br>Moder and Mor<br>humus | dystrophic to<br>mesotrophic /<br>moderate fresh /<br>moderate deep | T31 Temperate<br>mountain Picea<br>forest |

\* https://eunis.eea.europa.eu

### 2.1 Regional scale

The study focuses on three model regions with a high proportion of disturbed Norway spruce stands: The Western Thuringian Forest (TF), the Southern Harz Mountains (SH), and the Thuringian Slate Mountains (TS) (**Fig. 2**). According to the 2023 forest inventory by the forest authority (“ThüringenForst AöR”), the pre-disturbance shares of Norway spruce ranged from approximately 34% in the TF and SH regions to around 86% in TS. According to climatic classification based on records of the previous three decades, all three regions share a humid temperate climate, with annual precipitation sums accounting to 760 mm (SH), 880 mm (TF) and 914 mm (TS); annual mean temperatures range from 7.5 °C (TS) to 8.8 °C (TF) (Kronenberg et al., 2021). Cloud cover is frequent throughout the year, and snow cover may occur during mid-winter at higher elevations. The region “TF”, located in the western Thuringian Forest, lies within the South Thuringian Triassic hill landscape. The area is geologically diverse, comprising magmatic and metamorphic rocks (granite, gneiss), as well as sedimentary formations such as sandstone and conglomerate (TLUBN, 2006). The region “SH”, situated in the mid-elevation zones of the southern Harz Mountains, is characterized by geological substrates consisting of sedimentary rocks (greywacke, sandstone) and volcanic formations (rhyolite, andesite) (TLUBN, 2006). The region “TS” is located in the upper elevations of the Thuringian Slate Mountains and is geologically dominated by slate, along with other metamorphic and sedimentary rocks (sandstone, greywacke, quartzite, conglomerate) and volcanic rocks such as rhyolite (TLUBN, 2006). Across all model regions, the soils are predominantly cambisols, podzols, and leptosols, developed on acidic, silica-rich parent material. These geological substrates result in low to moderate nutrient availability, a characteristic shared across the regions. This edaphic context plays a crucial role in shaping forest regeneration dynamics following disturbance events. Post-disturbance management strategies of the forest offices differ notably across the three regions: in the region “TF”, the dominant approach has been small-scale salvage logging combined with the retention of standing dead trees. In contrast, the region “SH” exhibits a mixed management history, with disturbance areas encompassing both cleared and uncleared stands. In the region “TS”, by comparison, the predominant post-disturbance strategy has widely been the complete clearing of affected stands.

### 2.2 Local scale

The climatic and site conditions of individual disturbed areas may differ in characteristics such as local climate, water availability, soil properties, as well as deadwood and vegetation structure, all of which can reflect and affect the regeneration potential as well as the biodiversity of flora, fauna, and fungi. To document these individual characteristics and their variability, three study areas were established in each of the three model regions, each containing four plots of 1 ha in size. Each of the four treatment variants is therefore replicated three times within a model region, which results in a total of 36 plots of 1 ha each (3 model regions × 4 treatment variants × 3 replications; see **Fig. 2**). Experimental plots comprise different silvicultural strategies for dealing with disturbed areas at local scale, comprising the treatment variants D, S, C (**Fig. 3**), which are compared with undisturbed sites of Norway spruce as control (I).

**Figure 3:**
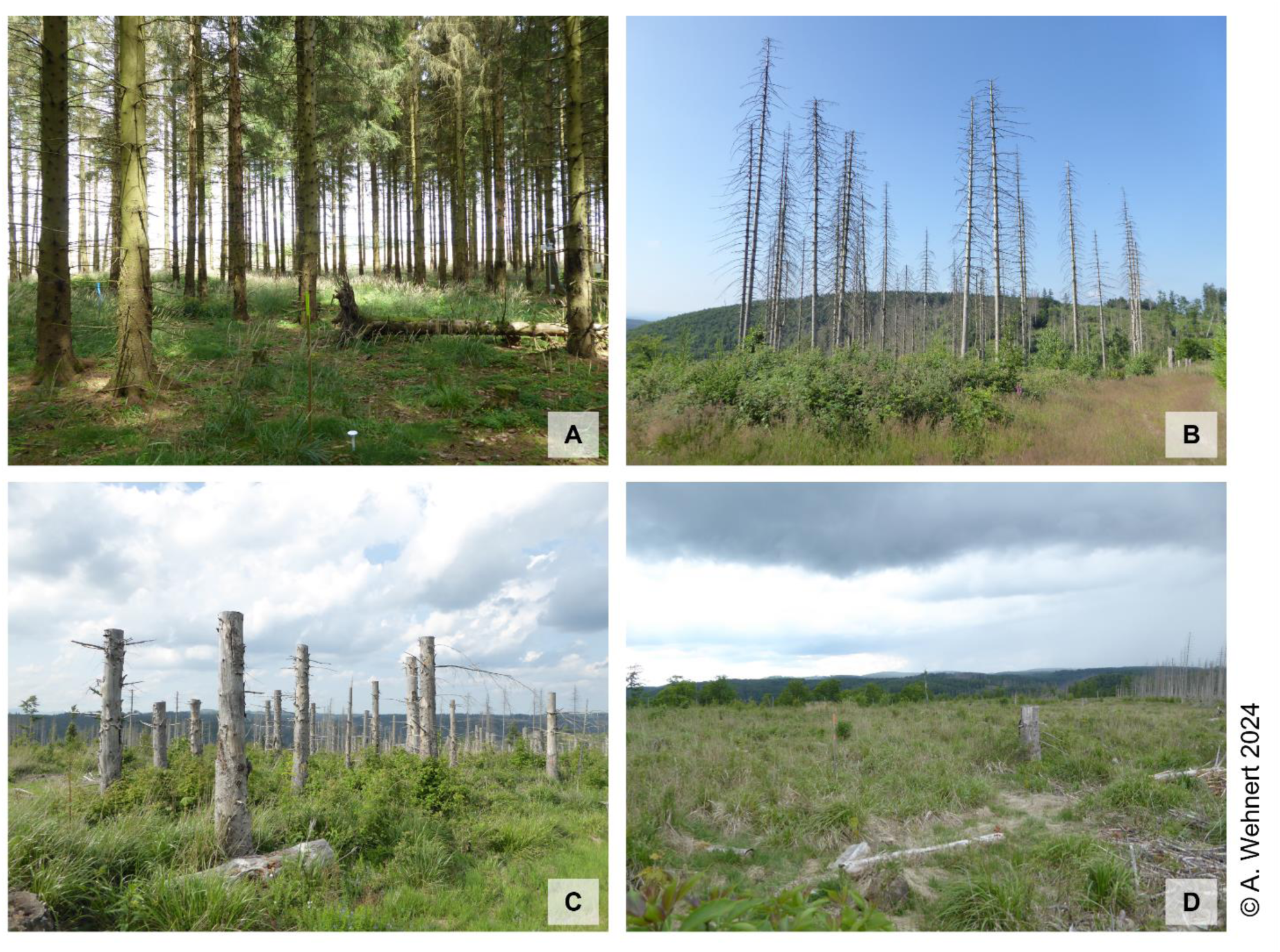
Exemplary photographs of the implemented variants for the post-disturbance deadwood retention treatment and undisturbed sites (1-ha-plots). A: undisturbed Norway spruce stand as experimental control for intact forest ‘I’ at study area Rothesütte in region SH; B: standing deadwood treatment ‘D’ at study area Etterwinden in region TF; C: high stumps treatment ‘S’; and D: clearing treatment ‘C’ at study area “Christianenhaus” in region SH.

#### 2.2.1 Implementation of plots for deadwood treatment

As working in areas of standing dead trees poses a significant health risk, the “standing deadwood” (D) treatment was implemented in the form of a checkerboard pattern of areas with dead trees, i.e. standing deadwood islands (Fig. 3B), and areas with removed dead trees. The latter offers higher safety (at least during times of low wind speed and previously stable weather conditions) and thus chances of long-term use of the installed measuring devices. Beyond their technical implementation benefits, standing deadwood islands also maintain larger areas free from soil compaction and mechanical disturbance, thereby providing refugia that support soil integrity and the occurrence of sensitive taxa such as fungi. Measurements are taken on subplots that are partly within and outside of standing dead trees, which makes them accessible for maintenance and monitoring while, e.g., the microclimate is still being affected by the shelter of the standing deadwood. For the treatment “high stumps” (S), dead trees were cut at approx. 2 m, leaving standing stumps. Occasionally, based on the decision of the local forest administration, individual stumps were left at greater height to provide perching opportunities for birds of prey. To implement the treatment option “clearing” (C), all dead tree trunks were roughly cut off at their root collars and coarse woody debris (CWD) was largely removed which left root stumps and trunk remnants with total heights of approx. 20-30 cm. In all treatments (D, S, C), fine woody debris (FWD) was left along the skid trails. Our assessment of all experimental plots after implementation of the treatment variants showed that the area covered by FWD differs between treatments as such: I <= C (0-20%) < S (20-40%) < D (40-60%).

Standing deadwood has been described as particularly valuable for ecological effects in relation to the natural decomposition process (Stokland, Siitonen, & Jonsson, 2012). However, when it comes to practical implementation of management concepts, leaving large areas with standing deadwood not only poses problems for work and forest infrastructure safety, but is also debated regarding its effect on recreation and tourism (Nielsen, Heyman, & Richnau, 2012; Rathmann, Sacher, Volkmann, & Mayer, 2020), depending on the location of the standing deadwood. As such, high stumps could be an easier-to-handle alternative to standing deadwood, preserving ecological value and less loss of timber value. To date, a holistic investigation on the effects of high stumps has not been performed, even though also small amounts of deadwood have been shown to provide microsites to foster tree establishment (Marzano et al., 2013; Steinebrunner et al., 2025; Swanson et al., 2023), for example in combination with plantings close to the old tree stumps (Schulze & Wagemann, 2020). A direct comparison between these practical deadwood variants can clarify how effects at different spatial levels and over time impact various ecological factors.

As implementation of treatments was carried out by local forest authorities individually, the amount of deadwood left on the subplots may differ depending on personal working experience, topography or machinery. Therefore, the deadwood elements with a diameter of more than 7 cm were recorded with coordinates within all 20 m x 20 m subplots. The decay classes and the volume of standing and lying deadwood were recorded based on the classification by Stokland (2001), Robin and Brang (2009), and Steventon (2011). Because disturbance dynamics differed among study areas, the treatment variants were implemented at different times. These implementation dates were documented and are considered in the analyses based on time since implementation.

#### 2.2.2 Implementation of regeneration subplots

Within each 1-ha-plot, two (I) or four (D, S, C) subplots of 20 m × 20 m each were implemented (**Fig. 2**), including two subplots (one for variant “I”) with natural regeneration (deadwood management without further intervention) and two subplots (one for variant “I”) with artificial regeneration (planting of Norway maple (*Acer platanoides* L.)). Compared to other tree species, also within the same genus, Norway maple is rarely applied in forestry practice. In contrast, Sycamore maple (*Acer pseudoplatanus* L.) is a mixed tree species of great ecological importance that is widespread in the low mountain regions (Caudullo & Rigo, 2016; Nowak & Rowntree, 1990). This is also reflected in the occurrence of seed trees and natural regeneration, but due to the increased occurrence of dry years and higher temperatures, Sycamore maples in particular are showing increased losses in vitality, which are further exacerbated by fungal infections (e.g. *Verticillium dahliae*, *Eutypella parasitica*, *Cryptostroma corticale*) (Burgdorf & Straßer, 2020). Because of its lower economic importance, Norway maple is less widespread and less is known about its robustness in the regeneration phase against the influencing factors mentioned (Carón et al., 2015). The ecological importance of Norway maple is considered to be high, especially on disturbed areas, which is due to the rapid juvenile growth, the favorable C/N ratio of the litter and the intensive rooting (Paquette et al., 2012; Roloff & Pietzarka, 2014). To test the establishment and growth potential of this species within the model regions, Norway maple was artificially planted on the respective subplots. The absence of seed trees for Norway maple also ensures that there is no mixing with the natural regeneration in the study areas. As a thermophilic pioneer tree species with low moisture requirements, this species is characterized by high drought tolerance and a broad physiological amplitude with regard to the pH values in the topsoil (Fuchs, 2021). Therefore, Norway maple is considered a suitable alternative to other pioneer tree species (e.g. *Sorbus aucuparia* L., *Salix caprea* L., *Populus tremula* L., *Betula pendula* Roth) for a rapid reforestation of disturbed areas. The subplots were not fenced against browsing to capture realistic browsing pressure, which represents a major limiting factor for natural regeneration in many Central European forests. This choice also aligns with our aim to co-evaluate management strategies under conditions representative of actual management contexts with foresters.

#### 2.2.3 Monitoring scheme

The subplots and their respective treatments are investigated through interdisciplinary monitoring to comprehensively assess climatic effects on soil, water, flora, fauna, and fungi. Taking this further, the fine-scale monitoring is then intended to be scaled up to the regional (landscape) level using UAV– and satellite-based remote sensing and long-term dynamics modeling to better understand how different management interventions influence ecosystem processes across different spatial and temporal scales. To ensure comparability across sites and disciplines and to capture interactions between local processes and regional dynamics, a continuous climatic data basis is fundamental for the characterization and evaluation of different management strategies.

In this context, it is important to distinguish between macroclimate and microclimate, as they capture different scales of variability and processes. Macroclimate generally refers to the broader atmospheric conditions measured in standardized settings (e.g., 2 m height above short, regularly cut grassland, following WMO guidelines), whereas microclimate describes the fine-scale climatic variation close to the ground or within vegetation canopies, strongly shaped by local topography, structure, and surface cover types (Geiger *et al*., 2009; Frenne *et al*., 2021). Distinguishing the two is relevant for our experimental approach because they are not equivalent in their ecological relevance: while macroclimatic measurements (e.g., air temperature at 2 m height) provide information on conditions affecting shrubs or more established trees, microclimatic variation reflects the spatial heterogeneity that seedlings, the herbaceous layer, or planted saplings are directly exposed to depending on surface cover types.

To account for the influence of different vegetation layers, each 1-ha plot in our study was equipped with a meteorological station to continuously record climatic conditions at plot-scale. The meteorological stations (model: µMETOS NB-IoT, Pessl Instruments) continuously (15-minute intervals) collect the following environmental variables on 36 research plots across the three model regions: global radiation, wind speed and direction (at 2 m height), precipitation, as well as air temperature and humidity (at 0.5 m and 2 m height). These measurements fall somewhat in between classical definitions: although they include variables typically considered macroclimatic, they are taken within forest stands where canopy height and openness substantially modify radiation, wind, and temperature regimes. Thus, they represent a plot-level, stand-modified macroclimate.

To record microclimatic conditions and spatial variability within the 1-ha plot, each subplot was additionally equipped with three soil moisture–temperature loggers (model: TMS4 Standard, TOMST). These data loggers measure at 15-minute intervals: soil temperature and moisture at –8 cm depth, as well as surface temperatures slightly above the ground (+2 cm) and at 15 cm above ground (+15 cm) (Wild et al., 2019).

## 3. Characterization of study areas

Despite a long history of repeated disturbances by snow breakage, windthrow, or bark beetle infestation in Norway spruce stands in the model regions, reforestation in the 19th and 20th centuries always took place with fast-growing Norway spruce trees due to economic considerations (Pinkard et al., 2015; Richter & Templin, 1980). Since the end of the 20^th^ century, there has been an increase in the frequency and extent of climate-related disturbances in the model regions (Senf & Seidl, 2018). Forest conversion measures are being continued on an ongoing basis in order to create climate-adapted forests (Apel et al., 2019; Frischbier et al., 2013). These forest conversion measures include enrichment with mixed tree species, promotion of natural regeneration via seed trees, use of adapted provenances, conversion of thinning measures to stabilize and structure the stands, and adjustment of game populations (Eichhorn, Guericke, & Eisenhauer, 2016; Frischbier, Profft, & Arenhövel, 2010; Fritz & Jenssen, 2006; Profft, Seiler, & Arenhövel, 2007), while the integration of deadwood into existing forest conversion concepts represents a new approach. Despite these measures in more places, their large-scale implementation and effects take time. Recent forest loss remains high as current climate extremes have reached unprecedented levels (Hartmann et al., 2025). The total volume of damaged timber accumulated up to August 2025 in the entire state of Thuringia amounts to approximately 23,250,000 solid cubic meters (fm). Within the Marksuhl Forestry District, the volume is approx. 232,000 fm; in the Bleicherode Forestry District (Southern Harz), about 436,000 fm; and in the Neuhaus Forestry District, approximately 2,194,000 fm (ThüringenForst AöR, 2025).

After disturbances, it is assumed that reforestation will begin through natural regeneration (Wohlgemuth, Jentsch, & Seidl, 2019). Different sequences in succession or targeted planting provide options for establishing multifunctional forests with varying structures that are better adapted to current and future environmental conditions and provide different ecosystem services. However, the speed of this development is strongly dependent on factors such as regional climate, distance to seed trees, accompanying vegetation, and others. Active intervention is or has often been carried out on disturbed areas to support reforestation through planting, but also to achieve the goal of changing tree species through forest conversion. It is therefore important when examining the effects of different deadwood management practices directly, to also consider local effects of the treatment variants, which are characterized by study area in **Table 1** and in the following. Detailed profiles of each study area can be found in the data publication by Profft and Spaleck (2025).

### 3.1 Economic aspects of implementing variants for deadwood treatment and regeneration

Many of the disturbed areas caused by drought and bark beetle infestation consist of standing deadwood, which can be left in place or harvested using conventional harvester operations. The use of harvesters and the removal of the wood incur costs for personnel and machinery, which depend on wage levels and the characteristics and accessibility of the area. The more impassable and remote the area, the higher the costs. Areas on steep slopes are unsuitable for the use of harvesters.

The quality of the wood from a dead tree may be reduced. Besides that, the value can be estimated in a similar way to felled trees. Key to the yield is the timber, which includes trunk wood and strong branch wood with a minimum diameter of 7 cm without bark. It is specified as merchantable wood volume in solid cubic meters and can further be roughly divided into the higher quality of saw timber (diameter > 14 cm) and industrial timber (diameter > 7 cm). Since the trunk of a tree progressively thins from bottom to top, there is no saw timber in the uppermost meters of the trunk. For small Norway spruce trees, this can be the top 8 of 14 m, for large trees the top 6 of 33 m (Pukkala, Holt Hanssen, & Andreassen, 2019). This consideration is important for comparing the yields that can be achieved for the high stump variant (S; wood below 2 m trunk height remains) compared to the complete clearing variant (C; cut at root collar) or standing deadwood (D; complete clearing of subplots, also with cut at root collar).

Let us consider, for example, the volume of a single, average Norway spruce tree with a diameter at breast height of 34 cm and a height of 22 m. For a rough estimate, we can use the volume formula that calculates the volume of a simple cylinder: V = f * g * H, where f = 0.5 is a correction factor for the taper of the trunk shape, g is the base area of the trunk cross-section at a height of 1.3 m, and H is the tree height. The so calculated volume of the example tree is 1 m³ (fm). If we compare this with a trunk that was cut at a height of 2 m and is 20 m high instead of 22 m and has a slightly reduced reference diameter of 32 cm instead of 34 cm at this height, we obtain a volume of 0.8 m³. The lower 9% of the trunk height therefore corresponds to approximately 20% of the trunk volume, which in the high stump variant does not contribute to the yield but remains on the ground as dead wood. Using taper functions that accurately describe the change in diameter with height in the trunk, the volume and relative differences can be calculated more precisely (e.g., Pukkala et al., 2019), but this general trend for larger trees remains: the relative loss represented by the lowest 2 m of the trunk is lower than for smaller trees (with the volume equation 35% for H = 20, DBH = 16 cm to 13% for H = 33, DBH = 50 cm). This is even more pronounced for the proportion of saw timber (> 14 cm diameter). In stands with small trees, the costs of setting up high stumps are therefore less likely to be offset by yields. More comprehensive analyses will show under which deadwood stand characteristics the high stump variant is advisable compared to leaving deadwood in place, and how cutting heights can be optimized for logs dependent on their diameter and height, i.e. striving for a minimum relative yield compared to complete removal.

For the establishment of standing deadwood, the comparison with clearing is easier because relative yield and relative costs appear to be directly proportional to the proportion of the area to be cleared around the deadwood islands. If 50% is cleared and the other 50% of the area remains as standing deadwood, this corresponds to 50% of the yield from complete clearing. Even if the costs for setting up standing deadwood do not correspond exactly to 50% of the costs of complete clearing, as spatial differentiation and longer travel distances make setting up standing deadwood somewhat less effective than complete clearing, it can still be assumed that this treatment is more cost-effective than setting up high stumps. However, with 50% of the area left as standing deadwood, the loss of yield is greater than the 13–35% resulting from the high stumps in the above calculation example. A more detailed breakdown of costs, timber stocks and sales prices will reveal the expected net proceeds of post-disturbance deadwood treatments for a portfolio of differently structured areas.

An economic assessment merely considering the short-term profit that can be achieved from the sale of deadwood is too short-sighted. There is also an economic value that arises in the medium to long term from the ecological impact of post-disturbance deadwood treatments. This includes ecosystem services that are difficult to quantify in monetary terms, such as the regulation of the water balance or the preservation of genetic diversity. And it can include effects that are directly relevant to forestry business calculations, e.g. regarding the success of artificial regeneration and the costs of reforestation, or regarding growth, resilience and average annual net yield.

To establish multifunctional forests on disturbed Norway spruce areas, the aim is to achieve a mix of tree species. This usually requires planting, which is costly. The extent of the disturbed areas also poses a challenge for forest companies and tree nurseries in terms of the provision of seed and planting material and its deployment. Where deciduous trees can still be found as seed trees, either directly on or in the vicinity of disturbed areas, the conversion into mixed forests is supported by natural processes of succession and forest development. Management that ensures the survival of these seed trees can benefit economically. In the study regions, the proportion of Norway spruce ranges from 34 to 86%, resulting in very different conditions for supporting natural regeneration on the disturbed areas. The necessary investment costs and challenges of implementing artificial regeneration on these sites therefore vary greatly. Added to this are the costs of protecting artificial regeneration from browsing, which depend on the necessary extent of planting measures and the local browsing pressure and hunting regime. Initial surveys in the subplots suggest that browsing pressure can vary greatly within a small area and that browsing of the very young regeneration a few years after disturbance seems to depend more on frequently used routes than on the characteristics of an area in terms of tree species and deadwood.

### 3.2 Initial post-disturbance status

#### 3.2.1 Tree species regeneration

The currently disturbed former Norway spruce stands are the result of extensive reforestation measures carried out after clear-cutting or large-scale disturbances in the past. After clear-cutting, trees were replanted at high densities per hectare, resulting in stands of the same age, single-layered, and homogeneous. Although the ecological importance of mixed tree species was well known, they were systematically removed from both old stands and regeneration areas for a long time (Deutsche Akademie der Landwirtschaftswissenschaften, 1962). The pure Norway spruce stands were often managed with high stem density and dense canopy closure, making natural regeneration rare. Due to these initial conditions, the current situation with large-scale disturbances is difficult for the establishment of natural regeneration. Depending on the size of the disturbed areas, only Norway spruce trees are present as seed trees in the surroundings of the disturbed areas, which is why Norway spruce trees dominate natural regeneration in most areas. Nevertheless, the area-specific regeneration densities are very heterogeneous, and large areas usually result in small-scale regeneration aggregates of seedlings, which are highly dependent on pre-regeneration, accompanying vegetation, and distance to the nearest seed trees.

The mean dispersal distances for Norway spruce seeds range between 35 m and 350 m (Gratzer & Waagepetersen, 2018; Piotti, Leonardi, Piovani, Scalfi, & Menozzi, 2009) so that spatial gradients in regeneration densities also develop due to climatic and light-related edge effects caused by old tree remains. The regeneration window on disturbed areas is also limited in terms of time for the establishment of early successional tree species, as the spread and competitive potential of grasses and *Rubus* ssp. species is very high. Early succession light-demanding tree genera (e.g., birch, poplar, willow, larch, pine) are known for their rapid colonization of areas but require a suitably favorable initial substrate such as vegetation-free or bare mineral soil areas. For these tree species, it is also the seed tree potential and the temporarily activated soil seed bank that are decisive (Tiebel, Huth, & Wagner, 2018). Due to the conditions already described in the former old stands, both prerequisites are unfavorable for high regeneration densities in most disturbed areas. Although the dispersal distances for wind-dispersed tree species are greater, spatially heterogeneous regeneration patterns also form in these tree species, providing clues as to where particularly good safe sites can be found on a small scale (Candotti, Ennemoser, Seeber, & Tomelleri, 2025; Schupp, 1995). Where mixed tree species can establish, intensive mixtures with strong interspecific competition during the regeneration phase can be expected in the future (Candotti et al., 2025).

#### 3.2.2 Characterization of herbs, shrubs and grasses

The pre-disturbance stands in all study areas were coniferous plantations of site-native trees, EUNIS type T3N (**Table 2**), thus the understory was typically species-poor and dominated by *Avenella flexuosa* (L. *Drejer*) and bryophyte species. In general, after a large-scale disturbance, early successional plant communities show a high diversity of perennial herb, grass, and shrub species (Swanson et al., 2011). The increased availability of light is the most relevant changing environmental factor. Diversity is additionally increased as forest understory species can persist after disturbance and grow alongside opportunistic or competitive species, such as *Rubus* spp. and *Calamagrostis* spp. (Swanson et al., 2011). After the initial phase, shrubs or grasses typically dominate the early successional plant community for several years before pioneer tree species outgrow them (Leuschner & Ellenberg, 2018). A comparable development can be observed in our plots. Given the predominant soil types in our study areas (**Table 1**), we expect the development of early successional plant communities typical of acidic or calcareous-poor sites (EUNIS R71 Willowherb and foxglove clearings) depending on the study location. A regional exception is the study area TS-2 in the Thuringian Slate Mountains, where species such as *Atropa bella-donna* (L.) and *Galeopsis tetrahit* (L.) that are typically part of calcareous-rich early successional plant communities (EUNIS R72 Burdock and deadly nightshade clearings) were observed during the 2024 vegetation period. We noticed that across all study areas, but particularly in the Thuringian Forest and parts of the Southern Harz, *Rubus idaeus* (L.) quickly became a dominant species following disturbance. The early successional vegetation in some study plots, for example, TS-1 and TS-3 in the Thuringian Slate Mountains, still includes species typical of the former forest stands in the short term, due to the short time since the disturbance and onset of management. Furthermore, early successional plant communities may overlap with species found in plant communities along forest fringes (EUNIS R52 Forest fringe of acidic nutrient-poor soils).

**Table 2:**
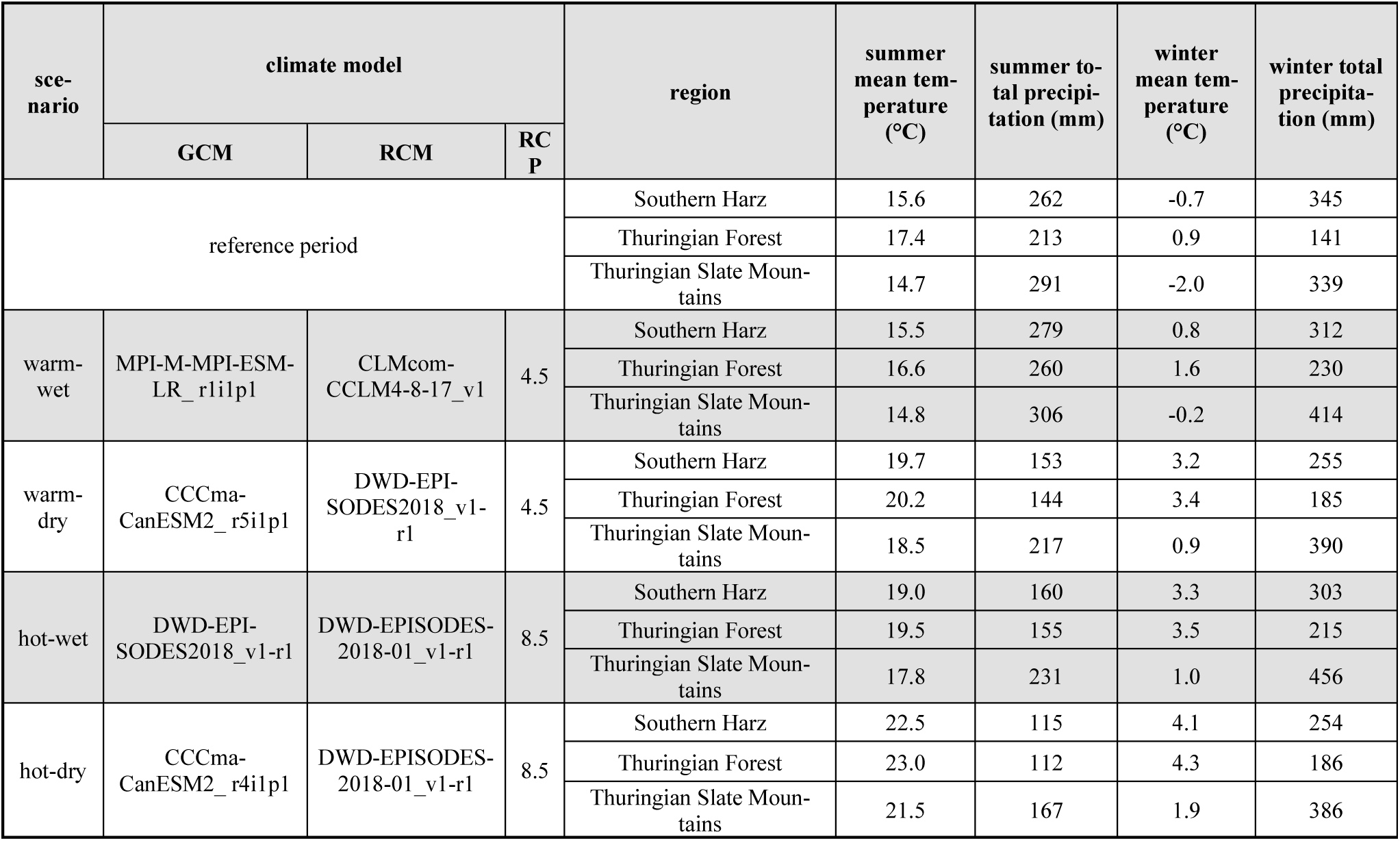
Summer (June–August) and winter (December–February) mean temperature and total precipitation for the three model regions: reference (1991–2020) from the nearest meteorological station with complete daily records, and scenario projections (2071–2100) at the representative coordinate in each region. The RCP (Representative Concentration Pathway) provides greenhouse gas/radiative forcing scenarios up to the year 2100. Given an RCP, a GCM (Global Climate Model) simulates global climate data, which is then refined at a regional level by an RCM (Regional Climate Model).

An important factor shaping the development and fate of juvenile trees and the early successional vegetation is the management of deadwood after disturbance. Previous studies showed that the removal of deadwood alters the early successional plant community (Steinebrunner et al., 2025). For example, salvage logging in bark-beetle-affected Lodgepole pine forests in North America increased species richness and grass cover but lowered shrub cover in the short term (Fornwalt et al., 2018). In Central European Norway spruce mountain forests, forest herbs survived under the dead canopy, whereas grasses dominated after salvage logging (Jonášová & Prach, 2008).

#### 3.2.3 Leaf area index and fraction of absorbed PAR

Before the different post-disturbance treatments may exhibit effects on vegetation development, the initial state of vegetation cover is affected by factors such as the type and frequency of forest machinery traffic, associated planting activities, the presence of surrounding disturbed areas or remaining trees and bushes contributing seed input, the duration of standing deadwood left after tree mortality before (complete or partial) removal, the timing of establishment (e.g., late autumn or spring), slight variations in the spacing of skid trails, and the amount and type of deadwood remaining on the plot, among others.

Against this background, we found that the initial condition of the study plots regarding vegetation status, characterized with biophysical parameters, differed considerably during the first vegetation period following the implementation of the management variant and planting. To characterize the variability in these initial conditions, *in-situ* measurements were carried out during peak vegetation season (June–July) using a hand-held ceptometer (model: SunScan, Delta-T Devices). Within the subplots, the mean Leaf Area Index (LAI) and the fraction of absorbed photosynthetically active radiation (FAPAR) were determined. For each subplot, the average of 20 individual measurements (one measurement per meter along a 20-meter transect) was calculated using the SunScan probe (1 m in length, equipped with 64 PAR sensors) oriented perpendicular to the transect. **Figure 4** shows the distribution of the mean LAI and mean FAPAR of the subplots on the disturbed site, covering all treatments. Most values (80^th^ percentile) fall below an LAI of 2.8 and a FAPAR of 0.8. The low LAI indicates an overall low to moderate vegetation cover. In contrast, LAI values of closed forest stands are considerably higher (e.g., >4 to 6). The FAPAR values also reflect the relatively open vegetation structure typical of disturbed or regenerating sites, suggesting bare areas and high to moderate solar radiation input. In sum, this suggests for an overall not yet closed, rather patchy vegetation cover on the investigated disturbed plots—with some variability, which is expected to have implications on microclimate, soil properties, succession rates and habitat function.

**Figure 4:**
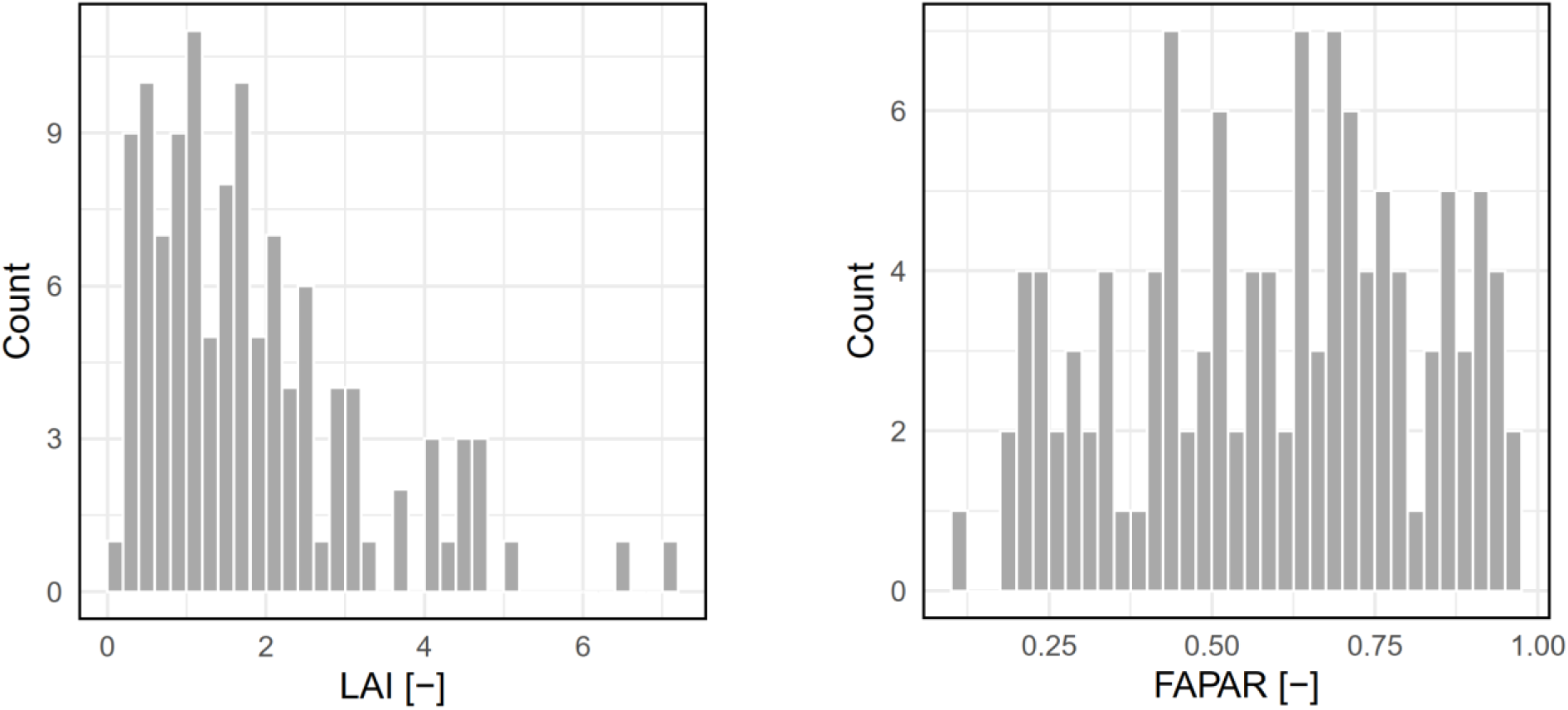
Leaf area index (LAI) and the fraction of actual photosynthetic active radiation (FAPAR) at the peak vegetation period after implementation of treatment variants (based on mean LAI in 117 subplots of treatment variants D, S, C). Note that treatment-specific differences were not depicted here, as potential effects are expected to emerge only after the initial establishment phase, while conditions in the first year primarily reflect site-specific and operational factors.

#### 3.2.4 Soil ecology and humus forms

Large-scale disturbances in Norway spruce forests lead to abrupt changes in ecological conditions of forest sites. The sudden loss of canopy alters litter input in quantitative and qualitative terms as well as temperature and moisture. These alterations induce changes in soil fauna and microbial abundance and activity, and thus in turn significantly affect humus form dynamics and soil ecological processes (Choma et al., 2021; Gomoryova et al., 2017; Salmon, Artuso, Frizzera, & Zampedri, 2008). In natural and managed Norway spruce stands, raw humus or moder forms typically dominate. These humus types are characterized by thick, acidic litter layers derived from poorly decomposed needle litter, high C:N ratios, and low biological activity and matter turnover. Limited faunal mixing promotes humic acid accumulation, podzolization, and weak nutrient release (Buresova et al., 2021; Zanella et al., 2018). Disturbance-induced shifts in vegetation and the cessation of needle litter input leads to dynamic changes in humus forms. These dynamics are closely linked to natural succession (Bernier & Ponge, 1994; Chauvat, Ponge, & Wolters, 2007). Pioneer species such as *Betula pendula*, *Sorbus aucuparia*, and grasses (e.g., *Calamagrostis* ssp., *Avenella flexuosa*) provide litter with higher quality compared to Norway spruce. Broadleaf and perennial litter is more decomposable and yields higher amounts of base cations, promoting transitions from raw humus to moder or mull forms (Klein-Raufhake et al., 2025; Labaz, Galka, Bogacz, Waroszewski, & Kabala, 2014). These processes are accompanied by the decreases in soil organic C by accelerated organic matter mineralization (Labaz et al., 2014), increases in soil pH following grass or broadleaf succession, altered N cycling, with enhanced nitrification and ammonification and shifts in microbial community composition (Choma et al., 2021). At the same time, soil fauna (especially earthworms and mesophilic invertebrates) can recolonize the ecosystems, enhancing litter fragmentation and incorporation in soil and humus mixing (Farská, Prejzková, Starý, & Rusek, 2014; Haimi & Huhta, 1990).

While these soil processes are largely governed by vegetation dynamics and abiotic conditions, forest management interventions, especially the extent to which deadwood is retained or removed, may critically influence soil ecological trajectories. Deadwood not only contributes both labile and recalcitrant organic substrates, but also regulates microclimate and provides habitat for saproxylic organisms, which may indirectly affect decomposition and nutrient cycling (Bani et al., 2018; Klein-Raufhake et al., 2025; Merganičová, Merganič, Svoboda, Bače, & ebe, 2012). To evaluate the suitability of different options of deadwood retention for sustainable forest management, their short and long-term effects on topsoil function, soil ecological status and turnover of organic matter will be assessed at the sub-plot scale within this project. Fine-scale humus layer sampling is combined with humus-form mapping to provide data that can be linked with successional dynamics.

#### 3.2.5 Microbial communities and related ecosystem processes

The effects of bark beetle-induced disturbances on Norway spruce stands vary depending on the lifestyle of the fungi involved. Ectomycorrhizal (ECM) fungi, which form mutualistic relationships with trees, generally experience declines in abundance and diversity following disturbances (Kosunen et al., 2020; Štursová et al., 2014; Veselá, Vašutová, Edwards-Jonášová, & Cudlín, 2019). A similar pattern was observed in our initial post-disturbance plots: ECM fungi accounted for around one-third of the fungal community in intact control plots (I), based on fruiting body inventories, and were almost exclusively associated with Norway spruce. This proportion was significantly lower in disturbed plots. For example, less than 5% of the fungal fruiting bodies in standing deadwood plots belonged to ECM species. However, an exception occurred in the Thuringian Forest (TF-1), where multiple ECM taxa associated with birch (*Betula pendula*) were present in clearing plots (C). The retention of ECM richness observed in these plots may be explained by the survival of a few local trees (Choma et al., 2023). The future course of ECM colonization depends on the successional pathways of tree regeneration (see Section 3.1.3). If Norway spruce dominates, diversity will mainly be recruited from the regional species pool. Colonization by other tree species, however, could broaden ECM associations. Regional contrasts would then likely play an important role because the Thuringian Forest’s higher pre-disturbance tree diversity provides a broader species pool for fungal recruitment than the Norway spruce-dominated Thuringian Slate Mountains do.

In contrast to ECM fungi, saprotrophic fungi typically benefit from the increased dead organic matter available after disturbance. Previous studies found that overall diversity increases, though the effects vary by substrate. Fungal communities in soil and litter showed initial declines in biomass and shifts in composition (Štursová et al., 2014), while deadwood exhibited the strongest changes in community structure (Masch, Buscot, Rohe, & Goldmann, 2025). At the start of our experiment, the amount of both coarse and fine woody debris differed strongly among treatments. Standing deadwood plots (D) had the highest amounts, followed by high stumps, clearings, and intact controls.

Since fungal diversity positively responds to substrate quantity and heterogeneity (Baber et al., 2016; Krah et al., 2018; Uhl et al., 2022), we expect both alpha and beta diversity to increase with greater deadwood availability over time. These effects may be further amplified as deadwood facilitates tree regeneration and new ECM hosts become established (Steinebrunner et al., 2025). Bacterial communities are generally more resistant to disturbance, with earlier studies reporting limited changes in species richness but shifts in community composition driven by altered soil conditions such as moisture, pH, and C:N ratio (Custer, van Diepen, & Stump, 2020; Štursová et al., 2014). A similar pattern may emerge in our experiment, with muted direct effects of treatments but indirect changes mediated by microclimate and nutrient availability.

#### 3.2.6 Bird and beetle diversity

The pre-disturbance stands in all study regions were pure Norway spruce stands, which supported a community of conifer-associated forest species, including coal tit (*Parus ater*) and goldcrest (*Regulus regulus*), as well as regionally rare species such as capercaillie (*Tetrao urogallus*) in the Thuringian Slate mountains region. Natural disturbances, such as bark beetle outbreaks, can significantly increase structural heterogeneity by altering light regimes and enhancing deadwood availability, thus creating habitat niches for disturbance-adapted specialist species (Thorn et al., 2016; Viljur et al., 2022). Structural elements remaining after disturbances, particularly standing deadwood and downed woody debris, are essential for conserving biodiversity, especially for organisms that rely on decaying wood for nutrition, reproduction, or shelter (Thorn et al., 2016; Thorn et al., 2020).

In this context, post-disturbance management treatments such as „standing deadwood” and “stumps” may offer new approaches to maintain or even enhance forest biodiversity. Retaining standing deadwood, for instance, can provide nesting habitats for cavity-nesting birds and offer temporary refugia for ground– and shrub-nesting species in open-canopy conditions (Castro, Moreno-Rueda, & Hódar, 2010). Leaving unlogged patches of standing deadwood within salvage-logged areas may thus support a more diverse bird assemblage during forest regeneration. For arthropods, canopy openness and the availability of deadwood are key drivers of diversity, particularly for saproxylic beetles (Perlík et al., 2023). While commonly neglected in post-disturbance management, the retention of high stumps can serve as important microhabitats and ecological steppingstones for wood-dependent species such as saproxylic beetles (Lackner, Reger, Tobisch, & Zahner, 2024). Given the recognized value of biological legacies in disturbed forests, stump retention may represent an underutilized tool in biodiversity-oriented management strategies. In addition, the natural regeneration of shrubs and herbaceous vegetation in post-disturbance forest sites provides critical resources for higher trophic levels.

### 3.3 Initial meteorological records in the year of planting

To assess the effects of intact Norway spruce forest cover on climatic conditions as a pre-disturbance baseline, we evaluated data from the aforementioned meteorological stations in the period of 01/11/2023 to 31/10/2024, deriving daily precipitation sum, daily radiation sum, and daily maximum (95^th^ percentile) 2-meter air temperature, and contrasting simultaneous measurements of the meteorological stations in cleared areas and intact stands within each study area (**Fig. 5**). Overall, a distinct influence of intact Norway spruce stands compared to clearings can be observed. Regarding precipitation, same-day sums are reduced for intact stands in most study areas, especially on days with high precipitation, with the slopes of a linear regression of daily precipitation sums between intact stands and clearings ranging from 0.32 to 0.91, thereby indicating the expected reduction in registered precipitation by interception within the intact stands. For daily radiation sums, the reduction in intact stands is most prominent (slopes ranging from 0.06 to 0.2), indicating a strong shading effect of Norway spruce trees compared to open, clear-cut areas. For 2-meter air temperature, there is a mitigation of temperature maxima in intact stands (slopes between 0.82 and 0.93). Overall, while the influence of intact stands is strongest for solar radiation, it is most consistent among study areas for the air temperature and has the highest variability for precipitation.

**Figure 5:**
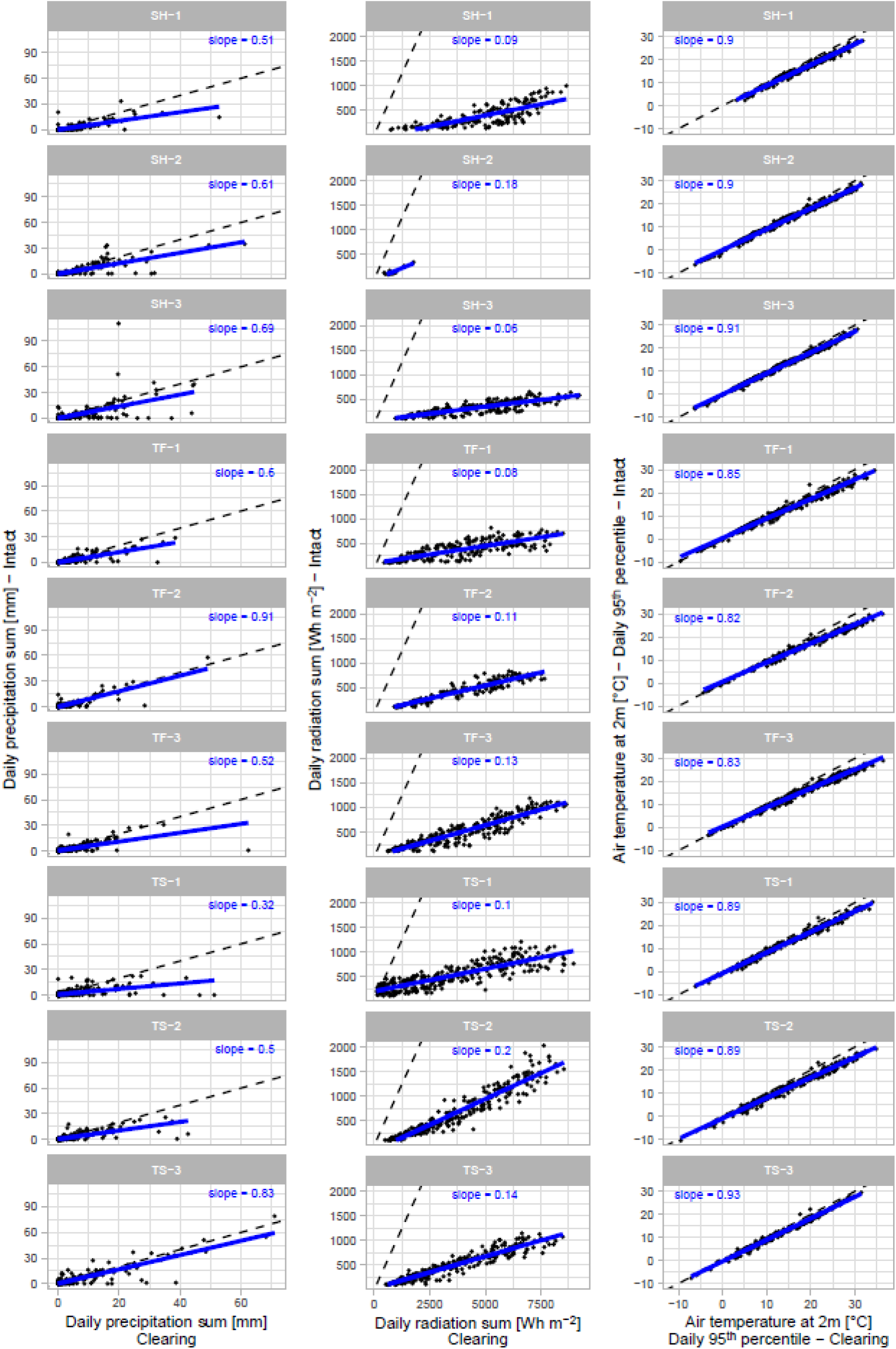
Daily precipitation sum, radiation sum and maximum temperature (95^th^ percentile) on cleared and intact plots with clearing vs. intact in each study area (01/11/2023-31/10/2024). Blue lines denote linear regressions, with the slope given for each study area. Dashed black lines are identity lines.

### 3.4 Predicted trends of climate change

To evaluate the potential impact of climate change on forest regeneration and subsequent long-term successional dynamics after disturbances in our study regions, we refer to climate projections from the “Mitteldeutsches Kernensemble” (MDK, v1.0; (Struve et al., 2020)), as provided by the ReKIS platform. MDK combines regionalized projections from EURO-CORDEX, ReKliEs-De and DWD EPISODES and, while maintaining the variability across climate variables, reduces the reference ensemble to seven model combinations per Representative Concentration Pathway scenario such as RCP4.5 and RCP8.5. The latter refer to rising greenhouse gas concentrations resulting in changes in radiative forcing of 4.5 and 8.5 W/m² in the year 2100 compared to the year 1750 (van Vuuren et al., 2011). We selected one representative coordinate for each of our study regions and downloaded the daily climate data for each point from ReKIS (scenario projections start in 2006). To cover the main range of temperature and precipitation changes, we used four MDK model combinations (**Fig. 6**). We label these combinations warm-wet and warm-dry under RCP4.5, and hot-wet and hot-dry under RCP8.5 for clarity (specific GCM, RCM and RCP of each combination are listed in **Table 2**). RCP4.5 takes the exhausting character of non-renewable fuels into account and is considered as the most probable baseline scenario, with emissions peaking around the year 2040. The RCP8.5 scenario represents a high-emission future pathway and is commonly utilized by regional forest authorities (e.g., ThüringenForst AöR) for forest and risk management planning. For the reference period (1991–2020), we used daily observations from the nearest meteorological station that had complete records for that period (sourced from ReKIS).

**Figure 6:**
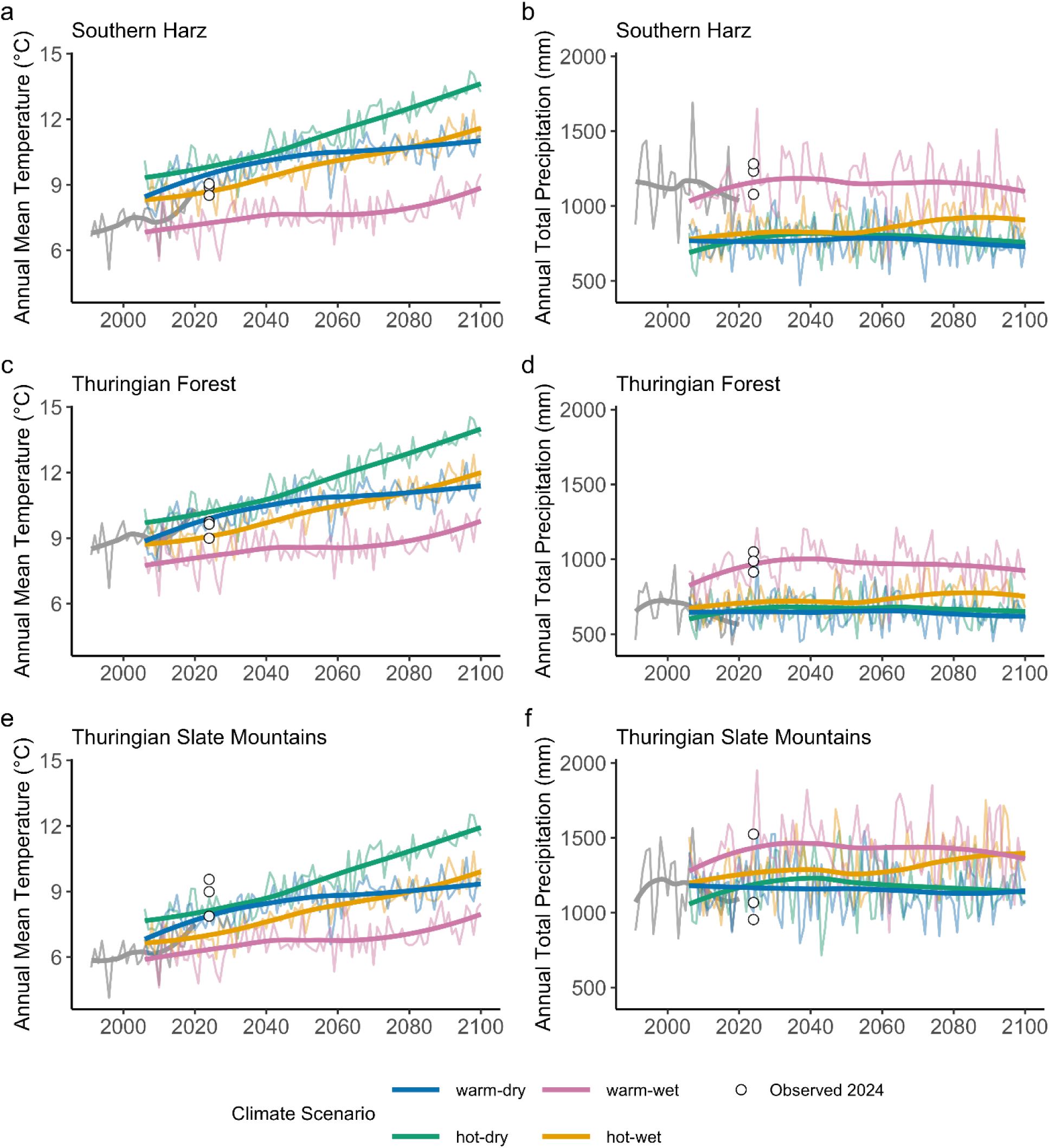
Projected annual mean temperatures (a, c, e) and total precipitation (b, d, f) under different climate scenarios at three representative coordinates (one per study region) from 2006 to 2100. Solid smooth lines represent LOESS smoothed trends for each scenario, while the thin jagged lines indicate annual data points. The grey lines depict observed data from the nearest meteorological stations during the reference period (1991–2020), while the scatter points represent measurements for the year 2024 obtained from the study areas (treatment variants C).

The projections indicate substantial warming trends across all three model regions by the end of the century. Under the most extreme hot-dry scenario, summerly mean temperatures are projected to increase by up to 6.9°C in Southern Harz, 5.6°C in Thuringian Forest, and 6.8°C in Thuringian Slate Mountains compared to the reference period. Winter temperatures show similar warming patterns, with increases ranging from 0.7°C to 4.8°C across regions and scenarios. Precipitation projections reveal distinct seasonal patterns with considerable regional variation. Summer precipitation is consistently projected to severely decrease across all except the ‘warm-wet’ scenario by 56 – 42% (Southern Harz), 48 – 27% (Thuringian Forest), 43 – 21% (Thuringian Slate Mountains). Conversely, winter precipitation is expected to increase substantially in most scenarios, with the largest increases projected under the warm-wet scenario in Thuringian Forest, exceeding 62%.

In summary, the climate model projections consistently point toward warmer temperatures, drier summers, and more humid winters across the model regions. These changes are expected to pose significant challenges to reforestation efforts, particularly during the critical summer growing season when water availability becomes limiting. By integrating these climate projections and the experimental framework into the individual-based forest landscape and disturbance model *iLand* (Rammer et al., 2024; Seidl, Rammer, Scheller, & Spies, 2012), simulations of long-term forest dynamics will be applied to evaluate how different deadwood management strategies and reforestation approaches perform under changing environmental conditions. Using such a simulation approach, the manifold empirical observations on soil properties, microclimate and regeneration, from the initial post-disturbance status to continued observations in treatment plots, can be assessed integrally and transferred into long term predictions of, i.e., forest growth, composition, structure and ecosystem functions such as timber production, carbon sequestration, site protection and biodiversity conservation. Besides performing simulations for different climate scenarios for empirically realized experimental treatments, additional simulations for synthetic experimental conditions will be carried out to reveal the effect of factors such as size of disturbance and seed tree availability, and how they translate into necessities and opportunities of post-disturbance management regarding the temporal and spatial aspects of planting.

## 4. Discussion

### 4.1 Representativeness and transferability across Central Europe

Forest disturbances in Central Europe have reached excessive magnitudes in severity, scale and frequency in the last decade and will remain a key challenge for forestry as climate change progresses. Already for the recent European drought years, critical thresholds for tree xylem hydraulic failure were reported to be crossed for many tree species, with strong drought legacy effects in consecutive years and a high vulnerability to secondary drought impacts such as insect or fungal infestation (Schuldt et al., 2020). Such vulnerability highlights the importance of understanding how site-specific conditions influence forest resilience.

To disentangle the effects of different environmental drivers on key response variables under study, cross-regional experimental designs are considered to be much more powerful than single region studies according to the orthogonality-representativeness-comprehensiveness concept (Verheyen et al., 2017). With multiregional data, repeated observations on the relationship between response variables to some environmental gradients are given, which allows inferences on the variability of such relationships and possible co-dependencies to other environmental factors. In this way, a cross-regional experimental design not only increases the ability to disentangle the effects of different environmental drivers and therefore creates higher orthogonality. It also generates a higher representativeness, as, simultaneously, a broader spectrum of environmental conditions is sampled and more reliable inference for a transferability of results is given. All of this also increases comprehensiveness, i.e. the spectrum of ecological questions that can be addressed with our experimental design. Despite these advantages of our cross-regional experimental design, our study focusses on specific natural areas.

The model regions in our study are representative of low mountain ranges in Central Europe, spanning from lower to mid and upper montane zones within their regions, with humid or partially very humid climate, and soils of low to moderate nutrient availability that developed on acidic, silica-rich parent material. The range of climate zones and levels of elevation allow for an upscaling of the results of this study to a broad range of areas in Central Europe that share these and other fundamental geopedological characteristics, for example, other parts of the German Central Upland and montane zones in the Bohemian Massif to the east and the Rhenish Massif to the west, in which summits usually do not reach the tree line and which were not glaciated in the last glacial period. However, transferability to otherwise very dry, base-rich sites or sites with deep soil and high nutrient supply or other sites in lowland areas is rather limited. Factors such as seed tree availability, seed bank, soil quality, and the composition of understory, microbial, and other species pools strongly depend on site history (section 3.1). Therefore, the transferability of our results to pre-disturbance stands other than ‘T3N coniferous plantations of site-native trees’ is limited. Nevertheless, as Norway spruce shares in the regions investigated range from 34% to 86%, our findings can still provide insights that are broadly applicable on the landscape scale. Finally, tree species related deadwood properties such as chemical structure, wood density and associated decaying rates and heat capacities can affect some linked ecological processes. As our field trial is based on Norway spruce only, additional research is needed to investigate the role of deadwood from other tree species, which is out of the scope of the current research approach. Apart from this critical view of possible limitations, the results of our study can be considered to contribute to key findings associated with the general response of forest systems to resource availability and microclimate, and possible dependencies of these responses on regional-specific environmental conditions, e.g. responses to microclimatic modifications due to deadwood treatment that depend on general macroclimatic conditions.

### 4.2 Climatic gradients under different post-disturbance management

Given these considerations on site characteristics and the potential transferability of our results, our experimental approach will provide a benchmark on how different post-disturbance management variants influence climatic conditions relevant for forest regeneration. Our first year of climatic observations already indicated distinct differences in temperature (extremes) between intact stands and open areas (**Fig. 5**). In clearings, higher solar radiation inputs led to a sharp rise in daytime air temperatures during the vegetation period, whereas intact forests effectively buffered such extremes, as common for forest interior climate (Frenne et al., 2021). Numerous studies have reported the distinct difference between closed canopy and open sites (Hofmeister et al., 2019; Kašpar et al., 2021). For the intermediate treatment variants “high stumps” and “standing deadwood”, we expect a gradient of increased buffering of temperature extremes. This expectation is supported by a recent three-year assessment of early post-disturbance conditions at one of our study areas (Putzenlechner et al., 2025), reporting distinct clear gradients across treatment variants: standing deadwood most effectively reduced summer temperature extremes, whereas high stumps did not differ significantly from the cleared plots which showed pronounced microclimatic amplification. Still, since only one hectare per variant was investigated, a holistic assessment across all study plots is now required to provide robust evidence of the microclimatic effects of the different management treatments. Nevertheless, these observations, together with existing studies on the role of microsites for forest regeneration (Lingua et al., 2023; Marzano et al., 2013; Steinebrunner et al., 2025), highlight the strong effect of structural legacies in modifying local microclimate. Whether such distinctions are detectable for air temperature measured at 2 m height remains uncertain, given the higher level of atmospheric mixing in this layer. In line with previous studies (Gril et al., 2023), we anticipate pronounced slopes of linear regression models between macroclimate, i.e. measurements in 2 m height, and microclimate, i.e. measurements near the soil surface, where structural elements and herbaceous vegetation can directly buffer radiation and evaporation (Atkins et al., 2023; Jucker et al., 2018; Zellweger et al., 2019).

In contrast to temperature, interpreting precipitation records will remain more challenging. The slopes of the linear regression models differed strongly depending on the specific location of the meteorological stations, which may also reflect the heterogeneous interception within stands. Differences between study areas could thus be explained not only by overall canopy openness of the intact stands but also by station placement relative to canopy gaps. Here, the additional observations from microclimatic dataloggers on soil moisture will provide valuable insights into spatial variability and improve the interpretation of treatment-specific differences.

From a temporal perspective, we expect microclimatic gradients to be most pronounced immediately following disturbance after implementation of the treatment variants, coinciding with a critical period for forest regeneration. This expectation is supported by initial findings from (Putzenlechner et al., 2025) who observed a homogenization of surface temperatures among treatments by the third year. However, over the long run, it is possible that, after such a plateau phase, the establishment of distinct plant communities will again generate differentiated microclimatic conditions across management variants over subsequent years. It is thus important to note that observable effects resulting from deadwood management depend on the time of observation and the speed of development, which is influenced by local (climatic) conditions. Our multi-year monitoring will allow us to capture these dynamics and provide insights into how structural legacies and vegetation development jointly shape local microclimates in the future (Máliš et al., 2023).

The climate projections indicate that summer droughts will occur more frequently and may even increase in severity in all three model regions (**Fig. 6**). This will challenge the drought resistance of both newly planted Norway maple trees and natural regeneration on our study plots as well as neighboring stands that have been less affected so far. This can in turn also influence seed dispersal to already disturbed sites. In this context, there is also uncertainty as to how long the plots with intact Norway spruce can sustain as reference. In our model regions, a high risk of further disturbances of remaining Norway spruce stands can be expected, which means that further disturbed areas would arise. As such, post-disturbance management strategies that help to buffer climate extremes would therefore be key to successful forest regeneration and conversion.

### 4.3 Trajectories of forest responses to drought

Despite the projected increase of drought frequency and severity, the response of forests observed in previous droughts may be different in the next sequence of drought events, as many vulnerable sites and trees have already been affected and resulted in changes in species composition and demographic structure, with associated modifications of drought vulnerability at these sites. Besides species– and age-specific responses to drought in the short-term, i.e. regarding stomatal closure (Hartmann, Link, & Schuldt, 2021), individual trees can develop different levels of drought resistance through phenotypic plasticity in allometry, leaf or xylem traits in response to experienced environmental conditions (Hacke, Spicer, Schreiber, & Plavcová, 2017; Kannenberg et al., 2022; Toca, Gonzalez-Benecke, Nelson, & Jacobs, 2025). This means that the survival of individual trees can result from an already acquired drought resistance and that regeneration, even of species that were previously severely affected, may develop less susceptible phenotypes at the same site and be able to withstand similar drought events in the future. However, our understanding of the species-specific rates and constraints of phenotypic plasticity is still limited. While the general drought resistance of Norway maple was recently ranked very high in a multi-criteria assessment (Leuschner, Fuchs, Wedde, Rüther, & Schuldt, 2024), indicating its suitability as a timber species in warmer and drier conditions, there are contrasting observations regarding the response of some species to natural or experimental drought. For Norway spruce, a high drought vulnerability is reflected by observed widespread disturbance rates in the last series of drought events (Putzenlechner *et al*., 2023). A generally higher drought vulnerability could, however, recently not be confirmed according to comparative analysis of inter-annual variations in physiological and leaf morphological traits (Schumann, Schuldt, Fischer, Ammer, & Leuschner, 2024). Similarly, Hesse et al. (2024) observed a successful mitigation of water stress by stomatal closure in the short term and reductions of the leaf area in the longer term in drought stress experiments with Norway spruce. Besides partly open questions on species-specific behavior, we are also just beginning to understand how the interaction of tree species can modify and partially improve their joint spatiotemporal water usage patterns (Hackmann et al., 2025). Moreover, the role of different types of deadwood retention strategies in the regeneration process was assessed differently (Leverkus et al., 2021a; Steinebrunner et al., 2025). Diversification in species and their genetic potential, as well as in silvicultural systems and management intensities in space and time (Huth et al., 2025), is to date considered as the most pragmatic strategy to strengthen forest resilience under uncertain future conditions and responses.

## 5. Perspective

The research approach presented here provides a multitude of indicators capturing macro– and microclimatic, soil chemical, biological, and structural responses to disturbance and management, providing an interdisciplinary perspective on the effects of disturbance management and deadwood retention strategies. In this regard, it will be investigated how the dynamics in humus forms and transformation processes are linked to disturbance and deadwood management as well as how these dynamics are linked to succession in ground vegetation and alterations in microclimate. Further, the experimental design will reveal how deadwood retention influences different microbial guilds as well as beetle and bird communities, and how these effects interact with vegetation succession across regions. Moreover, it will be clarified how microbial patterns and microclimatic variations among treatments translate into ecosystem processes such as decomposition and primary production via mutualistic interactions. Considering the spatiotemporal variability in abiotic and biotic conditions, the experimental setup and extensive monitoring including the recording of seed trees will provide a strong contribution to clarifying the manifold interactions affecting the initial seedling development and tree establishment. Linking these different disciplinary research efforts, our approach holds the potential to reveal how different levels of deadwood retention influence the near-surface microclimate and, in turn, how early-successional vegetation – such as grasses that may hinder tree regeneration – modulates soil conditions, feeding back to local (micro-)climatic dynamics, success of forest regeneration and tree growth.

The climatic measurements across different layers of the forest environment will provide a nuanced perspective that goes beyond the conventional distinction between micro– and macroclimate. While these categories are often used as separate concepts, the conditions we capture on our 1-ha plots likely fall somewhere in between, depending on canopy structure, vegetation height, and surface cover. This layered approach allows us to challenge and refine common definitions, showing that climatic conditions relevant for forest regeneration can vary along a gradient rather than fitting into one category. From an ecological perspective, such differentiation is particularly important in disturbed stands, where canopy removal alters climatic conditions at multiple levels in different ways. These conditions also change over time as succession proceeds, with vegetation growth gradually reshaping the near-ground environment. By simultaneously recording across different climatic layers, our design enables us to analyze how structural legacies and management treatments translate into regeneration-relevant environments – for seedlings and saplings close to the ground as well as for older cohorts exposed to stand-level conditions. Coupled with process-based modeling, this multi-layer perspective will provide a powerful tool to better understand regeneration pathways and to anticipate how management interventions may shape future forest development under changing climate conditions.

Upscaling will be realized by various approaches to optimally serve different purposes and dimensions (**Fig. 7**). Taking the results from the interdisciplinary assessment of treatment variants, recommendations for action for deadwood retention as part of a post-disturbance management, customizable for site conditions and specific forest development goals, will be developed. For this, upscaling will serve for a transfer of observed local effects and dynamics, e.g., of various ecosystem-specific services, to larger spatial scales, i.e., within a landscape or cross-regional, to an extended temporal scale, and finally to recommendations for action that allow forest managers to make science-informed decisions on the selection of management options or treatment plans for a customized support of ecosystem services.

**Figure 7:**
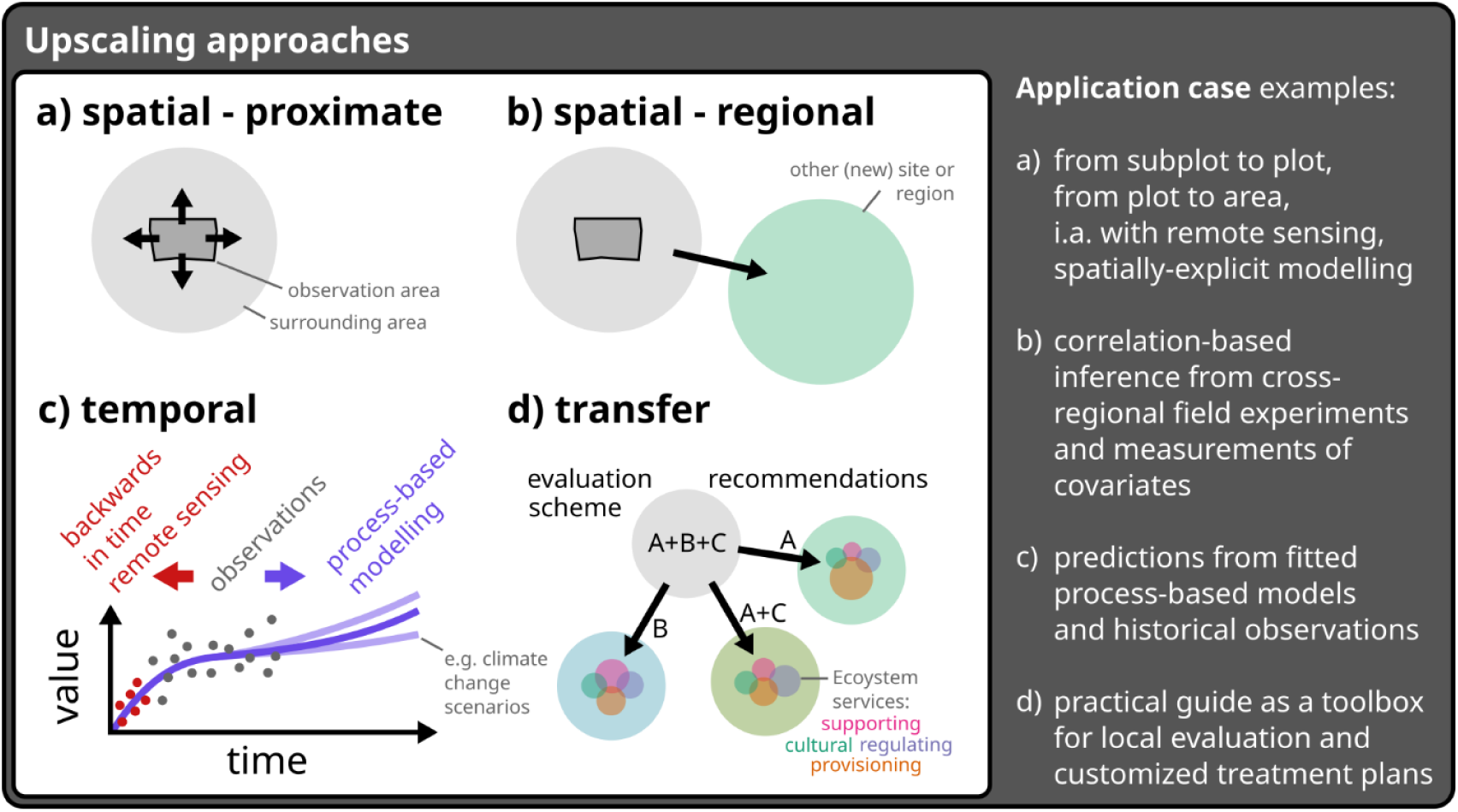
Upscaling approaches serving different dimensions and purposes, enabled by interdisciplinary research based on a cross-regional experimental setup. Upscaling can address the transfer of observed conditions and relationships to proximate, neighboring areas (a), to other regions (b), and across time (c). It can also address the transfer of recommendations to new scenarios of practical interest for forestry (d).

To achieve a spatio-temporal upscaling of results, we combine plot-scale observations with satellite– and UAV-based remote sensing, which not only enables assessment of vegetation recovery and microclimatic effects across broader regions but also provides a reconstruction of disturbance dynamics and insight into the current status quo. In addition, we use long-term forest dynamics modeling to explore potential future development of our and also of different management strategies, helping guide operational decisions under current uncertainty and supporting the development of monitoring approaches for forest regeneration at the landscape scale. As such, the cross-regional experimental design presented here allows us to capture spatial variability in multiple functions and begin tracking their temporal trajectories, which is essential for process-oriented analyses, upscaling of results, and transferability of recommendations to areas beyond our study plots. The plots set up for the ResEt-Fi project provide a valuable platform for additional research now and for future monitoring in order to resolve developmental trajectories that we are now beginning to track. We invite the scientific community to collaborate with us to harness and further develop this potential.

## Acknowledgements

This article is a joint initiative of the consortium of the “ResEt-Fi” collaborative project, funded by the Federal Ministry of Research, Technology and Space (BMFTR) (grant no. 033L304) as part of the REGULUS program (https://regulus-waldholz.de/en/ueber-regulus-regulus-waldholz-de/). The authors thank ThüringenForst AöR and the FFK Gotha as well as the local forest administrations for enabling and coordinating the research activities, providing and setting up forest stands for the field experiments.

## Author contributions

**BP**: Conceptualization, Resources, Data curation, Formal analysis, Investigation, Writing – Original Draft, Writing – Review & Editing, Supervision, Funding acquisition;

**MZ**: Conceptualization, Resources, Data curation, Formal analysis, Investigation, Writing – Original Draft, Writing – Review & Editing, Visualization, Supervision, Funding acquisition;

**CB**: Conceptualization, Resources, Writing – Review & Editing, Supervision, Funding acquisition;

**MB-R**: Conceptualization, Resources, Writing – Review & Editing, Supervision, Funding acquisition;

**JE**: Data curation, Investigation, Writing – Original Draft, Writing – Review & Editing;

**SG**: Formal analysis, Data curation, Investigation, Writing – Original Draft, Writing – Review & Editing, Visualization;

**PK**: Conceptualization, Data curation, Writing – Original Draft, Writing – Review & Editing, Funding acquisition, Visualization;

**HO**: Data curation, Investigation, Writing – Original Draft, Writing – Review & Editing;

**DP**: Data curation, Investigation, Writing – Original Draft, Writing – Review & Editing;

**IP**: Conceptualization, Resources, Writing – Review & Editing, Supervision, Project administration, Funding acquisition;

**SS**: Data curation, Investigation, Writing – Review & Editing;

**FS**: Data curation, Investigation, Writing – Original Draft, Writing – Review & Editing;

**AT**: Conceptualization, Resources, Writing – Review & Editing, Supervision, Funding acquisition;

**ST**: Conceptualization, Resources, Writing – Review & Editing, Supervision, Funding acquisition;

**AW**: Data curation, Investigation, Writing – Original Draft, Writing – Review & Editing;

**XZ**: Software, Data curation, Formal analysis, Investigation, Writing – Original Draft, Writing – Review & Editing, Visualization;

**FH**: Conceptualization, Data curation, Resources, Investigation, Writing – Original Draft, Writing – Review & Editing, Funding acquisition

## Data availability statement

Data on temperature, precipitation and radiation, recorded from 11/2023 to 10/2024 by the climate stations in the intact and clearing plots as well as data from LAI and FAPAR measurements is available upon reasonable request.

## Reference List

1. Albert, M., Nagel, R.-V., Nuske, R. S., Sutmöller, J., & Spellmann, H. (2017). Tree Species Selection in the Face of Drought Risk—Uncertainty in Forest Planning. Forests, 8(10).

2. Anders, T., Hetzer, J., Knapp, N., Forrest, M., Langan, L., Tölle, M. H., et al. (2025). Modelling past and future impacts of droughts on tree mortality and carbon storage in Norway spruce stands in Germany. Ecological Modelling, 501, 110987, from https://www.sciencedirect.com/science/article/pii/S0304380024003752.

3. Apel, J., Baier, U., Betzold, H., Burghoff, L., Dargel, H., Gehringer, M., et al. (2019). Forstchronik: die Geschichte des Waldes und der Forstwirtschaft in Thüringen: ThüringenForst.

4. Atkins, J. W., Shiklomanov, A., Mathes, K. C., Bond-Lamberty, B., & Gough, C. M. (2023). Effects of forest structural and compositional change on forest microclimates across a gradient of disturbance severity. Agricultural and Forest Meteorology, 339, 109566.

5. Baber, K., Otto, P., Kahl, T., Gossner, M. M., Wirth, C., Gminder, A., & Bässler, C. (2016). Disentangling the effects of forest-stand type and dead-wood origin of the early successional stage on the diversity of wood-inhabiting fungi. Forest Ecology and Management, 377, 161–169, from https://www.sciencedirect.com/science/article/pii/S0378112716303577.

6. Bani, A., Pioli, S., Ventura, M., Panzacchi, P., Borruso, L., Tognetti, R., et al. (2018). The role of microbial community in the decomposition of leaf litter and deadwood. Applied Soil Ecology, 126.

7. Bernier, N., & Ponge, J. F. (1994). Humus form dynamics during the sylvogenetic cycle in a mountain spruce forest. Soil Biology and Biochemistry, 26(2), 183–220, from https://www.sciencedirect.com/science/article/pii/0038071794901619.

8. Brockerhoff, E. G., Barbaro, L., Castagneyrol, B., Forrester, D. I., Gardiner, B., González-Olabarria, J. R., et al. (2017). Forest biodiversity, ecosystem functioning and the provision of ecosystem services. Biodiversity and Conservation, 26(13), 3005–3035.

9. Bundesministerium für Landwirtschaft, Ernährung und Heimat (BMLEH) (2025). Massive Schäden – Einsatz für Wälder und Waldumbau nötig. Retrieved July 23, 2025, from Bundesministerium für Landwirtschaft, Ernährung und Heimat (BMLEH): https://www.bmleh.de/DE/themen/wald/wald-in-deutschland/wald-trockenheit-klimawandel.html.

10. Buras, A., & Menzel, A. (2019). Projecting Tree Species Composition Changes of European Forests for 2061–2090 Under RCP 4.5 and RCP 8.5 Scenarios. Frontiers in Plant Science, 9(1986).

11. Buresova, A., Tejnecky, V., Kopecky, J., Drabek, O., Madrova, P., Rerichova, N., et al. (2021). Litter chemical quality and bacterial community structure influenced decomposition in acidic forest soil. European Journal of Soil Biology, 103, 103271, from https://www.sciencedirect.com/science/article/pii/S1164556320303411.

12. Burgdorf, N., & Straßer, L. (2020). Aktuelle pilzliche Erkrankungen bei Ahorn. Wald & Mehr, 1. Retrieved 05.08-2025, from https://www.lwf.bayern.de/waldschutz/phytopathologie/238091/index.php.

13. Candotti, A., Ennemoser, M., Seeber, J., & Tomelleri, E. (2025). Norway spruce dominates natural regeneration five years after a large-scale wind disturbance in the higher montane and lower subalpine belts in the eastern Alps. Forest Ecology and Management, 595, 123053, from https://www.sciencedirect.com/science/article/pii/S0378112725005614.

14. Carón, M. M., Frenne, P. de, Chabrerie, O., Cousins, S., Backer, L. de, Decocq, G., et al. (2015). Impacts of warming and changes in precipitation frequency on the regeneration of two Acer species. Flora – Morphology, Distribution, Functional Ecology of Plants, 214, 24–33, from https://www.sciencedirect.com/science/article/pii/S0367253015000456.

15. Castro, J., Moreno-Rueda, G., & Hódar, J. A. (2010). Experimental Test of Postfire Management in Pine Forests: Impact of Salvage Logging versus Partial Cutting and Nonintervention on Bird–Species Assemblages. Conservation Biology. (24), 810–819, from https://conbio.onlinelibrary.wiley.com/doi/abs/10.1111/j.1523-1739.2009.01382.x.

16. Castro, J., Allen, C. D., Molina-Morales, M., Marañón-Jiménez, S., Sánchez-Miranda, Á., & Zamora, R. (2011). Salvage Logging Versus the Use of Burnt Wood as a Nurse Object to Promote Post-Fire Tree Seedling Establishment. Restoration Ecology, 19(4), 537–544.

17. Caudullo, G., & Rigo, D. de (2016). Acer platanoides in Europe: distribution, habitat, usage and threats. In San-Miguel-Ayanz, J., Rigo, D., Caudullo, G., Houston Durrant, T., Mauri, A. (Ed.), European Atlas of Forest Tree Species. Luxembourg.

18. Chauvat, M., Ponge, J. F., & Wolters, V. (2007). Humus structure during a spruce forest rotation: quantitative changes and relationship to soil biota. European Journal of Soil Science, 58(3), 625–631.

19. Chianucci, F., Napoleone, F., Ricotta, C., Ferrara, C., Fusaro, L., Balducci, L., et al. (2024). Silvicultural regime shapes understory functional structure in European forests. Journal of Applied Ecology, 61(10), 2350–2364.

20. Choma, M., Bače, R., Čapek, P., Kaňa, J., Kaštovská, E., Tahovská, K., & Kopáček, J. (2023). Surviving trees are key elements in the fate of ectomycorrhizal community after severe bark-beetle forest disturbance. FEMS Microbiology Ecology, 99(8), fiad082.

21. Choma, M., Šamonil, P., Kaštovská, E., Bárta, J., Tahovská, K., Valtera, M., & Šantrůčková, H. (2021). Soil Microbiome Composition along the Natural Norway Spruce Forest Life Cycle. Forests, 12(4).

22. Custer, G. F., van Diepen, L. T. A., & Stump, W. L. (2020). Structural and Functional Dynamics of Soil Microbes following Spruce Beetle Infestation. Applied and Environmental Microbiology, 86(3), e01984–19.

23. Deutsche Akademie der Landwirtschaftswissenschaften (1962). Probleme der Waldökologie unter besonderer Berücksichtigung der Fichtenwirtschaft im Mittelgebirge: Vorträge anläßlich eines Symposiums vom 1. bis 7. Oktober 1961 im Institut für Forstwissenschaften Tharandt der Deutschen Akademie der Landwirtschaftswissenschaften zu Berlin. In Deutsche Akademie der Landwirtschaftswissenschaften (Ed.), Deutsche Akademie der Landwirtschaftswissenschaften zu Berlin*. Tagungsberichte* (Vol. 53).

24. Eichhorn, J., Guericke, M., & Eisenhauer, D.-R. (2016). Waldbauliche Klimaanpassung im regionalen Fokus. Sind unsere Wälder fit für den Klimawandel? Klimawandel in Regionen zukunftsfähig gestalten: Vol. 10. München, Germany: Ökom-Verlag.

25. Farská, J., Prejzková, K., Starý, J., & Rusek, J. (2014). Soil microarthropods in non-intervention montane spruce forest regenerating after bark-beetle outbreak. Ecological Research, 29, 1087–1096.

26. Fleischer, P., Pichler, V., Fleischer Jr, P., Holko, L., Máliš, F., Gömöryová, E., et al. (2017). Forest ecosystem services affected by natural disturbances, climate and land-use changes in the Tatra Mountains. Climate Research, 73(1-2), 57–71.

27. Fornwalt, P. J., Rhoades, C. C., Hubbard, R. M., Harris, R. L., Faist, A. M., & Bowman, W. D. (2018). Short-term understory plant community responses to salvage logging in beetle-affected lodgepole pine forests. Forest Ecology and Management, 409, 84–93, from https://www.sciencedirect.com/science/article/pii/S037811271731160X.

28. Forzieri, G., Girardello, M., Ceccherini, G., Spinoni, J., Feyen, L., Hartmann, H., et al. (2021). Emergent vulnerability to climate-driven disturbances in European forests. Nature communications, 12(1), 1081.

29. Frenne, P. de, Lenoir, J., Luoto, M., Scheffers, B. R., Zellweger, F., Aalto, J., et al. (2021). Forest microclimates and climate change: Importance, drivers and future research agenda. Global change biology, 27(11), 2279–2297.

30. Frischbier, N., Profft, I., & Arenhövel, W. (2010). Thuringian tree species recommendations for adaptation to climate change. Forst und Holz, 65, 28–35.

31. Frischbier, N., Profft, I., Richter, F., Feske, N., Stagl, J., Hattermann, F., & Gutsch, M. (2013). Habitat-change: Managing Spruce Forests Under Climate Change.

32. Fritz, P., & Jenssen, M. (2006). Ökologischer Waldumbau in Deutschland: Fragen, Antworten, Perspektiven: Ökom-Verlag.

33. Fuchs, S. M. (2021). *Diverse Forests for Climate Change: Drought Stress Tolerance of Secondary Timber Species.* Dissertation, Georg-August Universität, Göttingen, from https://ediss.uni-goettingen.de/handle/21.11130/00-1735-0000-0008-58E3-5.

34. Gohr, C., Blumröder, J. S., Sheil, D., & Ibisch, P. L. (2021). Quantifying the mitigation of temperature extremes by forests and wetlands in a temperate landscape. Ecological Informatics, 66, 101442.

35. Gomoryova, E., Fleischer, P., Pichler, V., Homolak, M., Gere, R., & Gomory, D. (2017). Soil microorganisms at the windthrow plots: the effect of post-disturbance management and the time since disturbance. iForest – Biogeosciences and Forestry, 10(2), 515–521, from https://iforest.sisef.org/contents/?id=ifor2304-010.

36. Gratzer, G., & Waagepetersen, R. P. (2018). Seed Dispersal, Microsites or Competition—What Drives Gap Regeneration in an Old-Growth Forest? An Application of Spatial Point Process Modelling. Forests, 9(5).

37. Grieger, S., Kappas, M., Karel, S., Koal, P., Koukal, T., Löw, M., et al. (2025). Impact of forest disturbance derived from Sentinel-2 time series on Landsat 8/9 land surface temperature: The case of Norway spruce in Central Germany. ISPRS Journal of Photogrammetry and Remote Sensing, 228, 388–407.

38. Gril, E., Spicher, F., Greiser, C., Ashcroft, M. B., Pincebourde, S., Durrieu, S., et al. (2023). Slope and equilibrium: A parsimonious and flexible approach to model microclimate. Methods in Ecology and Evolution, 14(3), 885–897.

39. Gustafsson, L., Baker, S. C., Bauhus, J., Beese, W. J., Brodie, A., Kouki, J., et al. (2012). Retention Forestry to Maintain Multifunctional Forests: A World Perspective. BioScience, 62(7), 633–645.

40. Hacke, U. G., Spicer, R., Schreiber, S. G., & Plavcová, L. (2017). An ecophysiological and developmental perspective on variation in vessel diameter. Plant, Cell & Environment, 40(6), 831–845.

41. Hackmann, C. A., Paligi, S. S., Mund, M., Hölscher, D., Leuschner, C., Pietig, K., & Ammer, C. (2025). Root water uptake depth in temperate forest trees: species-specific patterns shaped by neighbourhood and environment. Plant Biology.

42. Haimi, J., & Huhta, V. (1990). Effect of earthworms on decomposition processes in raw humus forest soil: A microcosm study. Biology and Fertility of Soils, 10(3), 178–183.

43. Hannerz, M., & Hånell, B. (1997). Effects on the flora in Norway spruce forests following clearcutting and shelterwood cutting. Forest Ecology and Management, 90(1), 29–49, from https://www.sciencedirect.com/science/article/pii/S0378112796038583.

44. Hartmann, H., Battisti, A., Brockerhoff, E. G., Bełka, M., Hurling, R., Jactel, H., et al. (2025). European forests are under increasing pressure from global change-driven invasions and accelerating epidemics by insects and diseases. Journal of Cultivated Plants, 77.

45. Hartmann, H., Link, R. M., & Schuldt, B. (2021). A whole-plant perspective of isohydry: stem-level support for leaf-level plant water regulation. Tree Physiol, 41(6), 901–905.

46. Hesse, B. D., Hikino, K., Gebhardt, T., Buchhart, C., Dervishi, V., Goisser, M., et al. (2024). Acclimation of mature spruce and beech to five years of repeated summer drought – The role of stomatal conductance and leaf area adjustment for water use. Science of The Total Environment, 951, 175805, from https://www.sciencedirect.com/science/article/pii/S0048969724059618.

47. Hesslerová, P., Huryna, H., Pokorný, J., & Procházka, J. (2018). The effect of forest disturbance on landscape temperature. Ecological Engineering, 120, 345–354.

48. Hlásny, T., König, L., Krokene, P., Lindner, M., Montagné-Huck, C., Müller, J., et al. (2021). Bark Beetle Outbreaks in Europe: State of Knowledge and Ways Forward for Management. Current Forestry Reports, 7(3), 138–165.

49. Hofmeister, J., Hošek, J., Brabec, M., Střalková, R., Mýlová, P., Bouda, M., et al. (2019). Microclimate edge effect in small fragments of temperate forests in the context of climate change. Forest Ecology and Management, 448, 48–56.

50. Huang, B., Li, Y., Liu, Y., Hu, X., Zhao, W., & Cherubini, F. (2023). A simplified multi-model statistical approach for predicting the effects of forest management on land surface temperature in Fennoscandia. Agricultural and Forest Meteorology, 332, 109362.

51. Huth, F., Tischer, A., Nikolova, P., Feldhaar, H., Wehnert, A., Hülsmann, L., et al. (2025). Ecological assessment of forest management approaches to develop resilient forests in the face of global change in Central Europe. Basic and Applied Ecology, 86, 66–100, from https://www.sciencedirect.com/science/article/pii/S1439179125000404.

52. Jonášová, M., & Prach, K. (2008). The influence of bark beetles outbreak vs. salvage logging on ground layer vegetation in Central European mountain spruce forests. Biological Conservation, 141(6), 1525–1535, from https://www.sciencedirect.com/science/article/pii/S0006320708001225.

53. Jucker, T., Hardwick, S. R., Both, S., Elias, D. M. O., Ewers, R. M., Milodowski, D. T., et al. (2018). Canopy structure and topography jointly constrain the microclimate of human-modified tropical landscapes. Global change biology, 24(11), 5243–5258.

54. Kannenberg, S. A., Guo, J. S., Novick, K. A., Anderegg, W. R. L., Feng, X., Kennedy, D., et al. (2022). Opportunities, challenges and pitfalls in characterizing plant water-use strategies. Functional Ecology, 36(1), 24–37.

55. Kašpar, V., Hederová, L., Macek, M., Müllerová, J., Prošek, J., Surový, P., et al. (2021). Temperature buffering in temperate forests: Comparing microclimate models based on ground measurements with active and passive remote sensing. Remote Sensing of Environment, 263, 112522, from https://www.sciencedirect.com/science/article/pii/S003442572100242X.

56. Keenan, R., & Kimmins, J. P. (1993). The ecological effects of clear-cutting. Environmental Reviews, 1(2), 121–144.

57. Klein-Raufhake, T., Hölzel, N., Schaper, J. J., Elmer, M., Fornfeist, M., Linnemann, B., et al. (2025). Disentangling the Impact of Forest Management Intensity Components on Soil Biological Processes. Global Change Biology, 31(1), e70018.

58. Knoke, T., Gosling, E., Thom, D., Chreptun, C., Rammig, A., & Seidl, R. (2021). Economic losses from natural disturbances in Norway spruce forests – A quantification using Monte-Carlo simulations. Ecological Economics, 185, 107046, from https://www.sciencedirect.com/science/article/pii/S092180092100104X.

59. Knutzen, F., Averbeck, P., Barrasso, C., Bouwer, L. M., Gardiner, B., Grünzweig, J. M., et al. (2025). Impacts on and damage to European forests from the 2018–2022 heat and drought events. Nat. Hazards Earth Syst. Sci., 25(1), 77–117, from https://nhess.copernicus.org/articles/25/77/2025/.

60. Kosunen, M., Peltoniemi, K., Pennanen, T., Lyytikäinen-Saarenmaa, P., Adamczyk, B., Fritze, H., et al. (2020). Storm and Ips typographus disturbance effects on carbon stocks, humus layer carbon fractions and microbial community composition in boreal Picea abies stands. Soil Biology and Biochemistry, 148, 107853, from https://www.sciencedirect.com/science/article/pii/S0038071720301504.

61. Krah, F.-S., Seibold, S., Brandl, R., Baldrian, P., Müller, J., & Bässler, C. (2018). Independent effects of host and environment on the diversity of wood-inhabiting fungi. The Journal of ecology, 106(4), 1428–1442.

62. Kronenberg, R., Franke, J., Neumann, T., Struve, S., Bernhofer, C., & Sommer, W. (2021). Das Regionale Klimainformationssystem ReKIS – eine gemeinsame Plattform für Sachsen, Sachsen-Anhalt und Thüringen. In P. Fischer-Stabel (Ed.), Umweltinformationssysteme. Grundlagen einer angewandten Geoinformatik/Geo-IT (3rd ed., pp. 1–9). Berling: Wichmann.

63. Kunert, N., Hajek, P., Hietz, P., Morris, H., Rosner, S., & Tholen, D. (2022). Summer temperatures reach the thermal tolerance threshold of photosynthetic decline in temperate conifers. *Plant biology (Stuttgart*, Germany), 24(7), 1254–1261.

64. Labaz, B., Galka, B., Bogacz, A., Waroszewski, J., & Kabala, C. (2014). Factors influencing humus forms and forest litter properties in the mid-mountains under temperate climate of southwestern Poland. Geoderma, *230-231*, 265–273, from https://www.sciencedirect.com/science/article/pii/S0016706114001785.

65. Lackner, T., Reger, B., Tobisch, C., & Zahner, V. (2024). The Potential of Artificial Snags to Promote Endangered Saproxylic Beetle Species in Bavarian Forests. Diversity, 16(5).

66. Laiho, R., & Prescott, C. E. (2004). Decay and nutrient dynamics of coarse woody debris in northern coniferous forests: a synthesis. Canadian Journal of Forest Research, 34(4), 763–777.

67. Leuschner, C., & Ellenberg, H. (2018). Ecology of Central European Forests: Vegetation Ecology of Central Europe, Volume I (1st ed.). Cham, Switzerland: Springer.

68. Leuschner, C., Fuchs, S., Wedde, P., Rüther, E., & Schuldt, B. (2024). A multi-criteria drought resistance assessment of temperate Acer, Carpinus, Fraxinus, Quercus, and Tilia species. Perspectives in Plant Ecology, Evolution and Systematics, 62, 125777, from https://www.sciencedirect.com/science/article/pii/S1433831923000616.

69. Leuschner, C. (2020). Drought response of European beech (Fagus sylvatica L.)—A review. Perspectives in Plant Ecology, Evolution and Systematics, 47, 125576.

70. Leverkus, A. B., Buma, B., Wagenbrenner, J., Burton, P. J., Lingua, E., Marzano, R., & Thorn, S. (2021a). Tamm review: Does salvage logging mitigate subsequent forest disturbances? Forest Ecology and Management, 481, 118721, from https://www.sciencedirect.com/science/article/pii/S0378112720314900.

71. Leverkus, A. B., Lindenmayer, D. B., Thorn, S., & Gustafsson, L. (2018). Salvage logging in the world’s forests: Interactions between natural disturbance and logging need recognition. Global Ecology and Biogeography, 27(10), 1140–1154.

72. Leverkus, A. B., Polo, I., Baudoux, C., Thorn, S., Gustafsson, L., & Rubio de Casas, R. (2021b). Resilience impacts of a secondary disturbance: Meta-analysis of salvage logging effects on tree regeneration. The Journal of ecology, 109(9), 3224–3232.

73. Lévesque, M., Saurer, M., Siegwolf, R., Eilmann, B., Brang, P., Bugmann, H., & Rigling, A. (2013). Drought response of five conifer species under contrasting water availability suggests high vulnerability of Norway spruce and European larch. Global change biology, 19(10), 3184–3199.

74. Lingua, E., Marques, G., Marchi, N., Garbarino, M., Marangon, D., Taccaliti, F., & Marzano, R. (2023). Post-Fire Restoration and Deadwood Management: Microsite Dynamics and Their Impact on Natural Regeneration. Forests, 14(9).

75. Máliš, F., Ujházy, K., Hederová, L., Ujházyová, M., Csölleová, L., Coomes, D. A., & Zellweger, F. (2023). Microclimate variation and recovery time in managed and old-growth temperate forests. Agricultural and Forest Meteorology, 342, 109722, from https://www.sciencedirect.com/science/article/pii/S0168192323004124.

76. Marangon, D., Marchi, N., & Lingua, E. (2022). Windthrown elements: a key point improving microsite amelioration and browsing protection to transplanted seedlings. Forest Ecology and Management, 508, 120050, from https://www.sciencedirect.com/science/article/pii/S0378112722000445.

77. Marshall, L. A. E., Fornwalt, P. J., Stevens-Rumann, C. S., Rodman, K. C., Rhoades, C. C., Zimlinghaus, K., et al. (2023). North-facing aspects, shade objects, and microtopographic depressions promote the survival and growth of tree seedlings planted after wildfire. Fire Ecology, 19(1), 26.

78. Marzano, R., Garbarino, M., Marcolin, E., Pividori, M., & Lingua, E. (2013). Deadwood anisotropic facilitation on seedling establishment after a stand-replacing wildfire in Aosta Valley (NW Italy). Ecological Engineering, 51, 117–122, from https://www.sciencedirect.com/science/article/pii/S0925857412003850.

79. Masch, D., Buscot, F., Rohe, W., & Goldmann, K. (2025). Bark beetle infestation alters mycobiomes in wood, litter, and soil associated with Norway spruce. FEMS Microbiology Ecology, 101(3), fiaf015.

80. Menge, J. H., Magdon, P., Wöllauer, S., & Ehbrecht, M. (2023). Impacts of forest management on stand and landscape-level microclimate heterogeneity of European beech forests. Landscape Ecology, 38(4), 903–917.

81. Merganičová, K., Merganič, J., Svoboda, M., Bače, R., & ebe, V. (2012). Deadwood in Forest Ecosystems. In J. H. Blanco & Y.-H. Lo (Eds.), Forest Ecosystems – More than Just Trees. London: IntechOpen.

82. Möhring, B., Bitter, A., Bub, G., Dieter, M., Dög, M., Hanewinkel, M., et al. (2021, March 05). Schadenssumme insgesamt 12,7 Mrd. Euro: Abschätzung der ökonomischen Schäden der Extremwetterereignisse der Jahre 2018 bis 2020 in der Forstwirtschaft. Holz-Zentralblatt, 9, 155–158, from https://www.waldwissen.net/assets/FVA/Waldwirtschaft/12_7_Mio_Schaden/2021_02_DFWR_Waldscha__den.pdf.

83. Morris, J. L., Cottrell, S., Fettig, C. J., DeRose, R. J., Mattor, K. M., Carter, V. A., et al. (2018). Bark beetles as agents of change in social–ecological systems. Frontiers in Ecology and the Environment, 16(S1), S34–S43.

84. Müller, J., Mitesser, O., Cadotte, M. W., van der Plas, F., Mori, A. S., Ammer, C., et al. (2023). Enhancing the structural diversity between forest patches—A concept and real-world experiment to study biodiversity, multifunctionality and forest resilience across spatial scales. Global Change Biology, 29(6), 1437–1450.

85. Nielsen, A. B., Heyman, E., & Richnau, G. (2012). Liked, disliked and unseen forest attributes: Relation to modes of viewing and cognitive constructs. Journal of environmental management, 113, 456–466, from https://www.sciencedirect.com/science/article/pii/S030147971200518X.

86. Nikinmaa, L., Koning, J. H. de, Derks, J., Grabska-Szwagrzyk, E., Konczal, A. A., Lindner, M., et al. (2024). The priorities in managing forest disturbances to enhance forest resilience: A comparison of a literature analysis and perceptions of forest professionals. Forest Policy and Economics, 158, 103119, from https://www.sciencedirect.com/science/article/pii/S1389934123002149.

87. Nirhamo, A., Hämäläinen, K., Junninen, K., & Kouki, J. (2023). Deadwood on clearcut sites during 20 years after harvests: The effects of tree retention level and prescribed burning. Forest Ecology and Management, 545.

88. Nowak, D. J., & Rowntree, R. A. (1990). History and Range of Norway maple. Journal of Arboriculture. (16), 291–296.

89. Nyland, R. D. (2016). Silviculture: Concepts and Applications*, Third Edition:* Waveland Press.

90. Obladen, N., Dechering, P., Skiadaresis, G., Tegel, W., Keßler, J., Höllerl, S., et al. (2021). Tree mortality of European beech and Norway spruce induced by 2018-2019 hot droughts in central Germany. Agricultural and Forest Meteorology, 307, 108482, from https://www.sciencedirect.com/science/article/pii/S0168192321001659.

91. Paquette, A., Fontaine, B., Berninger, F., Dubois, K., Lechowicz, M. J., Messier, C., et al. (2012). Norway maple displays greater seasonal growth and phenotypic plasticity to light than native sugar maple. Tree Physiology, 32(11), 1339–1347.

92. Patacca, M., Lindner, M., Lucas-Borja, M. E., Cordonnier, T., Fidej, G., Gardiner, B., et al. (2023). Significant increase in natural disturbance impacts on European forests since 1950. Global change biology, 29(5), 1359–1376.

93. Perlík, M., Kraus, D., Bußler, H., Neudam, L., Pietsch, S., Mergner, U., et al. (2023). Canopy openness as the main driver of aculeate Hymenoptera and saproxylic beetle diversity following natural disturbances and salvage logging. Forest Ecology and Management, 540, 121033, from https://www.sciencedirect.com/science/article/pii/S0378112723002670.

94. Pinkard, E., Battaglia, M., Bruce, J., Matthews, S., Callister, A. N., Hetherington, S., et al. (2015). A history of forestry management responses to climatic variability and their current relevance for developing climate change adaptation strategies. Forestry, 88(2), 155–171.

95. Piotti, A., Leonardi, S., Piovani, P., Scalfi, M., & Menozzi, P. (2009). Spruce colonization at treeline: where do those seeds come from? Heredity, 103(2), 136–145.

96. Profft, I., Seiler, M., & Arenhövel, W. (2007). Die Zukunft der Fichte in Thüringen vor dem Hintergrund des Klimawandels. Forst und Holz. (62), 19–25.

97. Profft, I. & Spaleck, S. (2025). Characteristics of the study areas for post-disturbance deadwood management project “ResEt-Fi, from https://zenodo.org/records/17587138.

98. Puettmann, K. J., Wilson, S. M., Baker, S. C., Donoso, P. J., Drössler, L., Amente, G., et al. (2015). Silvicultural alternatives to conventional even-aged forest management – what limits global adoption? Forest Ecosystems, 2(1), 8.

99. Pukkala, T., Holt Hanssen, K., & Andreassen, K. (2019). Stem taper and bark functions for Norway spruce in Norway. Silva Fennica, 53(3), from https://silvafennica.fi/article/10187.

100. Putzenlechner, B., Grieger, S., Czech, C., & Koal, P. (2025). Post-disturbance treatment effects on microclimate and vegetation recovery on Norway spruce calamity areas from in situ and UAV-based monitoring. Forest Ecology and Management, 597, 123131, from https://www.sciencedirect.com/science/article/pii/S0378112725006395.

101. Rammer, W., Thom, D., Baumann, M., Braziunas, K., Dollinger, C., Kerber, J., et al. (2024). The individual-based forest landscape and disturbance model iLand: Overview, progress, and outlook. Ecological Modelling, 495, 110785, from https://www.sciencedirect.com/science/article/pii/S030438002400173X.

102. Rathmann, J., Sacher, P., Volkmann, N., & Mayer, M. (2020). Using the visitor-employed photography method to analyse deadwood perceptions of forest visitors: a case study from Bavarian Forest National Park, Germany. European Journal of Forest Research, 139(3), 431–442.

103. Richter, D., & Templin, E. (1980). Forstschutzsituation und –maßnahmen unter besonderer Berücksichtigung der Schnee-und Sturmschadgebiete (No. 8), pp. 240–241.

104. Robin, V., & Brang, P. (2009). Erhebungsmethode für liegendes Totholz in Kernflächen von Naturwaldreservaten. Eidg. Forschungsanstalt für Wald, Schnee und Landschaft (WSL).

105. Roloff, A., & Pietzarka, U. (2014). Acer platanoides. In Enzyklopädie der Holzgewächse: Handbuch und Atlas der Dendrologie (pp. 1–16).

106. Salmon, S., Artuso, N., Frizzera, L., & Zampedri, R. (2008). Relationships between soil fauna communities and humus forms: Response to forest dynamics and solar radiation. Soil Biology and Biochemistry, 40, 1707–1715.

107. Schroeder, L. M. (2007). Retention or salvage logging of standing trees killed by the spruce bark beetle Ips typographus: Consequences for dead wood dynamics and biodiversity. Scandinavian Journal of Forest Research, 22(6), 524–530.

108. Schuldt, B., Buras, A., Arend, M., Vitasse, Y., Beierkuhnlein, C., Damm, A., et al. (2020). A first assessment of the impact of the extreme 2018 summer drought on Central European forests. Basic and Applied Ecology, 45, 86–103, from https://www.sciencedirect.com/science/article/pii/S1439179120300414.

109. Schulze, L. & Wagemann, O. (2020). Mutterstöcke und Stockachselpflanzung zur Wiederbewaldung von Kalamitätsflächen, from https://www.waldwissen.net/de/waldwirtschaft/waldbau/mutterstoecke-und-stockachselpflanzung-zur-wiederbewaldung-von-kalamitaetsflaechen.

110. Schumann, K., Schuldt, B., Fischer, M., Ammer, C., & Leuschner, C. (2024). Xylem safety in relation to the stringency of plant water potential regulation of European beech, Norway spruce, and Douglas-fir trees during severe drought. Trees, 38(3), 607–623.

111. Schupp, E. W. (1995). Seed-seedling conflicts, habitat choice, and patterns of plant recruitment. American Journal of Botany, 82(3), 399–409.

112. Seibold, S., Bässler, C., Brandl, R., Büche, B., Szallies, A., Thorn, S., et al. (2016). Microclimate and habitat heterogeneity as the major drivers of beetle diversity in dead wood. Journal of Applied Ecology, 53(3), 934–943.

113. Seidl, R., Rammer, W., Scheller, R. M., & Spies, T. A. (2012). An individual-based process model to simulate landscape-scale forest ecosystem dynamics. Ecological Modelling, 231, 87–100, from https://www.sciencedirect.com/science/article/pii/S0304380012000919.

114. Senf, C., Buras, A., Zang, C. S., Rammig, A., & Seidl, R. (2020). Excess forest mortality is consistently linked to drought across Europe. Nature communications, 11(1), 6200.

115. Senf, C., & Seidl, R. (2018). Natural disturbances are spatially diverse but temporally synchronized across temperate forest landscapes in Europe. Global change biology, 24(3), 1201–1211.

116. Senf, C., & Seidl, R. (2021). Storm and fire disturbances in Europe: Distribution and trends. Global change biology, 27(15), 3605–3619.

117. Spiecker, H., Hansen, J., Klimo, E., Skovsgaard, J. P., Sterba, H., & Teuffel, K. von (2004). Norway Spruce Conversion: Options and Consequences. Leiden, Niederlande: Brill.

118. Steinebrunner, F., Tischer, A., Medicus, T., Huth, F., & Bernhardt-Römermann, M. (2025). The effects of deadwood on tree regeneration and microsites: A systematic review. Forest Ecology and Management, 596, 123096, from https://www.sciencedirect.com/science/article/pii/S0378112725006048.

119. Steventon, J. D. (2011). Retention patches: Windthrow and recruitment of habitat structure 12–16 years after harvest. Journal of Ecosystems and Management. (11), 18–28, from http://jem.forrex.org/index.php/jem/article/view/18/36.

120. Stokland, J. N. (2001). The Coarse Woody Debris Profile: An Archive of Recent Forest History and an Important Biodiversity Indicator. Ecological Bulletins. (49), 71–83, from http://www.jstor.org/stable/20113265.

121. Stokland, J. N., Siitonen, J., & Jonsson, B. G. (2012). Biodiversity in Dead Wood. Ecology, Biodiversity and Conservation. Cambridge: Cambridge University Press.

122. Storch, I., Penner, J., Asbeck, T., Basile, M., Bauhus, J., Braunisch, V., et al. (2020). Evaluating the effectiveness of retention forestry to enhance biodiversity in production forests of Central Europe using an interdisciplinary, multi-scale approach. Ecology and Evolution, 10(3), 1489–1509.

123. Struve, S., Ehlert, I., Pfannschmidt, K., Heyner, F., Franke, J., Kronenberg, R., & Eichhorn, M. (2020). Mitteldeutsches Kernensemble zur Auswertung regionaler Klimamodelldaten – Dokumentation – Version 1.0. Halle (Saale): Landesamt für Umweltschutz Sachsen-Anhalt (LAU), from https://rekis.hydro.tu-dresden.de/wp-content/uploads/2020/05/Dokumentation_Mitteldeutsches_Kernensemble_MDK.pdf.

124. Štursová, M., Šnajdr, J., Cajthaml, T., Bárta, J., Šantrůčková, H., & Baldrian, P. (2014). When the forest dies: the response of forest soil fungi to a bark beetle-induced tree dieback. The ISME Journal, 8(9), 1920–1931.

125. Swanson, M., Magee, M., Nelson, A., Engstrom, R., & Adams, H. (2023). Experimental downed woody debris-created microsites enhance tree survival and growth in extreme summer heat. Frontiers in Forests and Global Change, 6.

126. Swanson, M. E., Franklin, J. F., Beschta, R. L., Crisafulli, C. M., DellaSala, D. A., Hutto, R. L., et al. (2011). The forgotten stage of forest succession: early-successional ecosystems on forest sites. Frontiers in Ecology and the Environment, 9(2), 117–125.

127. Thonfeld, F., Gessner, U., Holzwarth, S., Kriese, J., Da Ponte, E., Huth, J., & Kuenzer, C. (2022). A First Assessment of Canopy Cover Loss in Germany’s Forests after the 2018–2020 Drought Years. Remote Sensing, 14(3), 562.

128. Thorn, S., Bässler, C., Bernhardt-Römermann, M., Cadotte, M., Heibl, C., Schäfer, H., et al. (2016). Changes in the dominant assembly mechanism drive species loss caused by declining resources. Ecology Letters, 19(2), 163–170.

129. Thorn, S., Bässler, C., Brandl, R., Burton, P. J., Cahall, R., Campbell, J. L., et al. (2018). Impacts of salvage logging on biodiversity: a meta-analysis. Journal of Applied Ecology, 55(1), 279–289.

130. Thorn, S., Seibold, S., Leverkus, A. B., Michler, T., Müller, J., Noss, R. F., et al. (2020). The living dead: acknowledging life after tree death to stop forest degradation. Frontiers in Ecology and the Environment, 18(9), 505–512.

131. ThüringenForst AöR (2023). Waldschutz-Information Nr. 4/2023: Befallssituation Buchdrucker und Kupferstecher. Gotha: ThüringenForst AöR. Retrieved August 27, 2025, from https://www.thueringenforst.de/fileadmin/user_upload/PDF/Waldschutzinformation/Waldschutzinformation_2023_04-Buchdrucker.pdf.

132. ThüringenForst AöR (2025). Waldschutz-Information Nr. 5/2025: Buchdrucker-Situation in Thüringen. Gotha: ThüringenForst AöR. Retrieved August 27, 2025, from https://www.thueringenforst.de/fileadmin/user_upload/PDF/Waldschutzinformation/Waldschutzinformation-5-2025-Buchdrucker.pdf.

133. Tiebel, K., Huth, F., & Wagner, S. (2018). Soil seed banks of pioneer tree species in European temperate forests: a review. iForest – Biogeosciences and Forestry, 11(1), 48–57, from https://iforest.sisef.org/contents/?id=ifor2400-011.

134. TLUBN (2006). Bodengeologische Karte von Thüringen / Soil geological map of Thuringia. Weimar: Thüringer Landesamt für Umwelt, Bergbau und Naturschutz (TLUBN).

135. Toca, A., Gonzalez-Benecke, C. A., Nelson, A. S., & Jacobs, D. F. (2025). Drought memory expression varies across ecologically contrasting forest tree species. Environmental and Experimental Botany, 231, 106094, from https://www.sciencedirect.com/science/article/pii/S0098847225000115.

136. Uhl, B., Krah, F.-S., Baldrian, P., Brandl, R., Hagge, J., Müller, J., et al. (2022). Snags, logs, stumps, and microclimate as tools optimizing deadwood enrichment for forest biodiversity. Biological Conservation, 270, 109569, from https://www.sciencedirect.com/science/article/pii/S0006320722001227.

137. van Vuuren, D. P., Edmonds, J., Kainuma, M., Riahi, K., Thomson, A., Hibbard, K., et al. (2011). The representative concentration pathways: an overview. Climatic Change, 109(1-2), 5–31.

138. Verheyen, K., Frenne, P. de, Baeten, L., Waller, D. M., Hédl, R., Perring, M. P., et al. (2017). Combining Biodiversity Resurveys across Regions to Advance Global Change Research. BioScience, 67(1), 73–83.

139. Veselá, P., Vašutová, M., Edwards-Jonášová, M., & Cudlín, P. (2019). Soil Fungal Community in Norway Spruce Forests under Bark Beetle Attack. *Forests,* 10(2).

140. Viljur, M.-L., Abella, S. R., Adámek, M., Alencar, J. B. R., Barber, N. A., Beudert, B., et al. (2022). The effect of natural disturbances on forest biodiversity: an ecological synthesis. Biological Reviews, 97(5), 1930–1947.

141. West, E., Morley P. J., Sump, A. S., & Donoghue, D. N. M. (2022). Satellite data track spatial and temporal declines in European beech forest canopy characteristics associated with intense drought events in the Rhön Biosphere Reserve, central Germany. Plant Biology. (24), 1120–1131. Retrieved May 10, 2023, from 10.1111/plb.13391.

142. Wild, J., Kopecký, M., Macek, M., Šanda, M., Jankovec, J., & Haase, T. (2019). Climate at ecologically relevant scales: A new temperature and soil moisture logger for long-term microclimate measurement. Agricultural and Forest Meteorology, 268, 40–47.

143. Wohlgemuth, T., Jentsch, A., & Seidl, R. (2019). Störungsökologie. (1. Auflage, Ed.): Haupt Verlag.

144. Yachi, S., & Loreau, M. (1999). Biodiversity and ecosystem productivity in a fluctuating environment: the insurance hypothesis. Proceedings of the National Academy of Sciences, 96(4), 1463–1468.

145. Zanella, A., Ponge, J. F., Jabiol, B., Sartori, G., Kolb, E., Le Bayon, R. C., et al. (2018). Terrestrial humus systems and forms – keys of classification of humus systems and forms. Applied Soil Ecology, 122(Part 1), 75–86.

146. Zellweger, F., Coomes, D., Lenoir, J., Depauw, L., Maes, S. L., Wulf, M., et al. (2019). Seasonal drivers of understorey temperature buffering in temperate deciduous forests across Europe. Global Ecology and Biogeography, 28(12), 1774–1786.

147. Zhang, S., Sjögren, J., & Jönsson, M. (2024). Retention forestry amplifies microclimate buffering in boreal forests. Agricultural and Forest Meteorology, 350, 109973.

148. Zimmermann, B., Kruber, S., Nendel, C., Munack, H., & Hildmann, C. (2024). Assessing the cooling potential of climate change adaptation measures in rural areas. Journal of environmental management, 366, 121595, from https://www.sciencedirect.com/science/article/pii/S0301479724015810.

